# Nde1 is Required for Heterochromatin Compaction and Stability in Neocortical Neurons

**DOI:** 10.1101/2021.06.25.449848

**Authors:** Alison A. Chomiak, Clara C. Lowe, Yan Guo, Dennis McDaniel, Hongna Pan, Xiaoming Zhou, Qiong Zhou, Martin L. Doughty, Yuanyi Feng

## Abstract

The *NDE1* gene encodes a scaffold protein essential for brain development. While biallelic *NDE1* loss of function (LOF) causes microcephaly with profound mental retardation, *NDE1* missense mutations and copy number variations are associated with multiple neuropsychiatric disorders. However, the etiology of the diverse phenotypes resulting from *NDE1* aberrations remains elusive. Here we show Nde1 controls neurogenesis through facilitating heterochromatin compaction via histone H4K20 trimethylation. This mechanism patterns diverse chromatin landscapes and stabilizes constitutive heterochromatin of neocortical neurons. We show NDE1 undergoes dynamic liquid-liquid phase separation, partitioning to the nucleus and interacting with pericentromeric and centromeric satellite repeats. Nde1 LOF results in nuclear architecture aberrations and DNA double strand breaks, as well as instability and derepression of pericentromeric satellite repeats in neocortical neurons. These findings uncover a pivotal role of NDE1/Nde1 in establishing and maintaining neuronal heterochromatin. They suggest that heterochromatin impairments underlie a wide range of brain dysfunction.

## INTRODUCTION

Although the expansion of cerebrum size is one of the most discernible characteristics of mammalian evolution, the increased number of cortical neurons is not sufficient to account for the increased intellectual capacities of the human brain. Along with this notion, genetic conditions that affect the generation of cortical neurons may lead to similar reductions in brain size but with distinctive cognitive outcomes. In contrast to some microcephaly (small brain) conditions with preserved cognition (Rauch et al., 2008; Rossi et al., 1987; Willems et al., 2010), biallelic loss of function (LOF) mutations of *NDE1* (NudE neurodevelopment protein 1) cause profound mental retardation (Alkuraya et al., 2011; Bakircioglu et al., 2011; Guven et al., 2012; Paciorkowski et al., 2013). While the near-completely diminished brain function resulting from severe NDE1 deficiency is attributed to the substantial neuronal loss, changes in brain size and morphology are not observed in individuals carrying missense mutations of *NDE1* or copy number variations (CNVs) of the *NDE1* gene locus 16p13.11. However, these genetic variants of *NDE1* are associated with a diverse array of neuropsychiatric disorders, including intellectual disability, epilepsy, autism, schizophrenia, and attention-deficit hyperactive disorder (Allach El Khattabi et al., 2020; Hannes et al., 2009; Ingason et al., 2011; Kimura et al., 2015; Nagamani et al., 2011; Ramalingam et al., 2011; Tropeano et al., 2013; Ullmann et al., 2007; Williams et al., 2010). The wide spectrum of brain morphological and functional impairments associated with *NDE1* gene dosage and function aberrations indicates that NDE1 acts beyond simply determining the quantity of neurons. However, the cellular and molecular mechanisms that underlie the diverse brain functional deficits resulting from *NDE1* LOF remain undefined.

The *NDE1* gene encodes a dynamic scaffold molecule that can be partitioned into various subcellular compartments in a context-dependent manner (Feng et al., 2000; Feng and Walsh, 2004; Lanctot et al., 2013; Pawlisz and Feng, 2011). Notably, the phenotype of NDE1/Nde1 LOF in both humans and mice preferentially affects the central nervous system (CNS), showing a prominent impact on neocortical neurogenesis (Alkuraya et al., 2011; Bakircioglu et al., 2011; Feng and Walsh, 2004; Guven et al., 2012). Despite the marked loss of neurons of the neocortex, homozygous Nde1 knockout (Nde1^-/-^) mice showed little anatomical change in other brain structures such as the hippocampus and cerebellum (Feng and Walsh, 2004). We have shown previously that the neocortical-specific neuronal loss was a direct consequence of Trp53-dependent apoptosis, which was triggered by DNA double-strand breaks (DSBs) in those Nde1^-/-^ neural progenitor cells (NPCs) that underwent the transition from multipotent to neuronal-fate-restrictive differentiations at the onset of cortical neurogenesis. More specifically, DSBs in Nde1^-/-^ NPCs arose from stalled replication of heterochromatin DNA in mid to late S-phase (Houlihan and Feng, 2014). The establishment of heterochromatin regions is fundamental to stem cell differentiation, and it reshapes the nuclear landscape to allow the spatial segregation of chromatin domains with distinctive transcription activities (Craig, 2005; Noma et al., 2001; Sridharan et al., 2013; Talbert and Henikoff, 2006). Therefore, this finding suggested that the pivotal role of NDE1 in neurogenesis is associated with the necessary heterochromatin remodeling for neuronal differentiation.

Heterochromatin constitutes a large portion of the mammalian genome (Lander et al., 2001; Linhoff et al., 2015; Solovei et al., 2016; Vicient and Casacuberta, 2017). It is distinguished from the transcriptionally active and loosely packed euchromatin by forming condensed structures that are predominantly distributed near the nuclear periphery and the nucleoli. While the formation of heterochromatin can be facultative, driven by developmental- and/or cell type-specific epigenetic transcriptional repression programs, the chromatin segments composed of tandem repeats of the centromere, pericentromere, subtelomere, and transposons must be silenced constitutively to maintain chromosome integrity and to prevent the genetic instability of these repetitive DNA elements (Allshire and Madhani, 2018; Saksouk et al., 2015; Vicient and Casacuberta, 2017). Despite a crucial role in genome maintenance and function, constitutive heterochromatin is the most poorly understood component of the eukaryotic genome, owing to difficulties in assessing its underlying repetitive DNA structures and the lack of genetic experimental models to study its in vivo formation, regulation, and function.

Given the specific requirement of Nde1 in linking heterochromatin replication and genome protection to neuronal differentiation, we asked whether the multitude of phenotypes resulting from *NDE1* aberrations originates from aberrant heterochromatin establishment, maintenance, stability, and/or function. In this study, we present evidence that Nde1 LOF impairs trimethylation of histone H4 lysine 20 (H4K20me3), a histone posttranslational modification (PTM) that distinctively marks the constitutive heterochromatin and stabilizes the repetitive genomic sequences. We found that H4K20me3 is not only required for neurogenesis but also shapes diverse chromatin landscapes of cortical neurons in coordination with brain development, aging, and evolution. We also show that Nde1 is an intrinsically disordered protein (IDP), can undergo liquid-liquid phase separation, and is directed by its partner, Brap, to the nucleus, where Nde1 interacts with the centromeric and pericenteromeric heterochromatin. Furthermore, mice with LOF of Brap phenocopy Nde1^-/-^ mice. Both show not only altered constitutive heterochromatization during cortical neurogenesis, but also heterochromatin instability in neocortical neurons, resulting in DSBs, profound nuclear architecture aberrations, and de-repression of pericentromeric satellite repeats. These heterochromatin defects coincided with progeria-like changes as well as neuroinflammation and neurodegenerative phenotypes in the cerebral cortical tissue, which are likely to underlie the cognitive dysfunction and multitude of neuropsychiatric phenotypes associated with *NDE1* aberrations.

## RESULTS

### Cerebral Cortical Neurogenesis Requires Heterochromatin Compaction via H4K20me3

To test whether the role of NDE1 in cerebral cortex neurogenesis is associated with re-patterning heterochromatin in NPC differentiation, we first determined how heterochromatin is modified in neurogenesis. We analyzed PTMs on histones extracted from mouse embryonic cortical tissues over the course of cerebral cortex development by immunoblotting (IB). From the onset of cortical neurogenesis at embryonic day 12.5 (E12.5) to its completion at birth (P0), we found that the levels of histone H4 lysine 20 trimethylation (H4K20me3), histone H3 lysine 27 trimethylation (H3K27me3), and histone H2A ubiquitination (H2Aub) increased progressively in the cortical tissue (Figure 1A). The increases in H3K27me3 and H2Aub were expected for their roles in the suppression of polycomb-group genes and the establishment of facultative heterochromatin in neurodevelopment. However, the remarkable upregulation of H4K20me3 and corresponding down regulation of H4K20me1, was surprising. H4K20me3 is a repressive histone modification enriched at the repetitive DNA elements of constitutive heterochromatin that are condensed in early stem cell differentiation and are expected to be stable throughout the cells’ lifespan (Benetti et al., 2007; Jorgensen et al., 2013; Kidder et al., 2017; Saksouk et al., 2015; Schotta et al., 2004; Schotta et al., 2008). The de novo establishment of constitutive heterochromatin requires epigenetic silencing by histone H3 lysine 9 trimethylation (H3K9me3), which is followed by H4K20me3 to further condense repetitive DNA elements and transposons to ensure their stability (Bierhoff et al., 2014; Jorgensen et al., 2013; Marion et al., 2009; Richards and Elgin, 2002; Schotta et al., 2008; Wongtawan et al., 2011). Consistent with this, we observed that the intensity of H3K9me3 foci was uniform in all cortical cells, but the intensity of H4K20me3 foci drastically increased in new neurons (DCX+ or Tuj1+) compared to NPCs, even in those newborn neurons migrating from the ventricular zone to the cortical plate (Figure 1B-D, Figure S1A).

**Figure 1.**
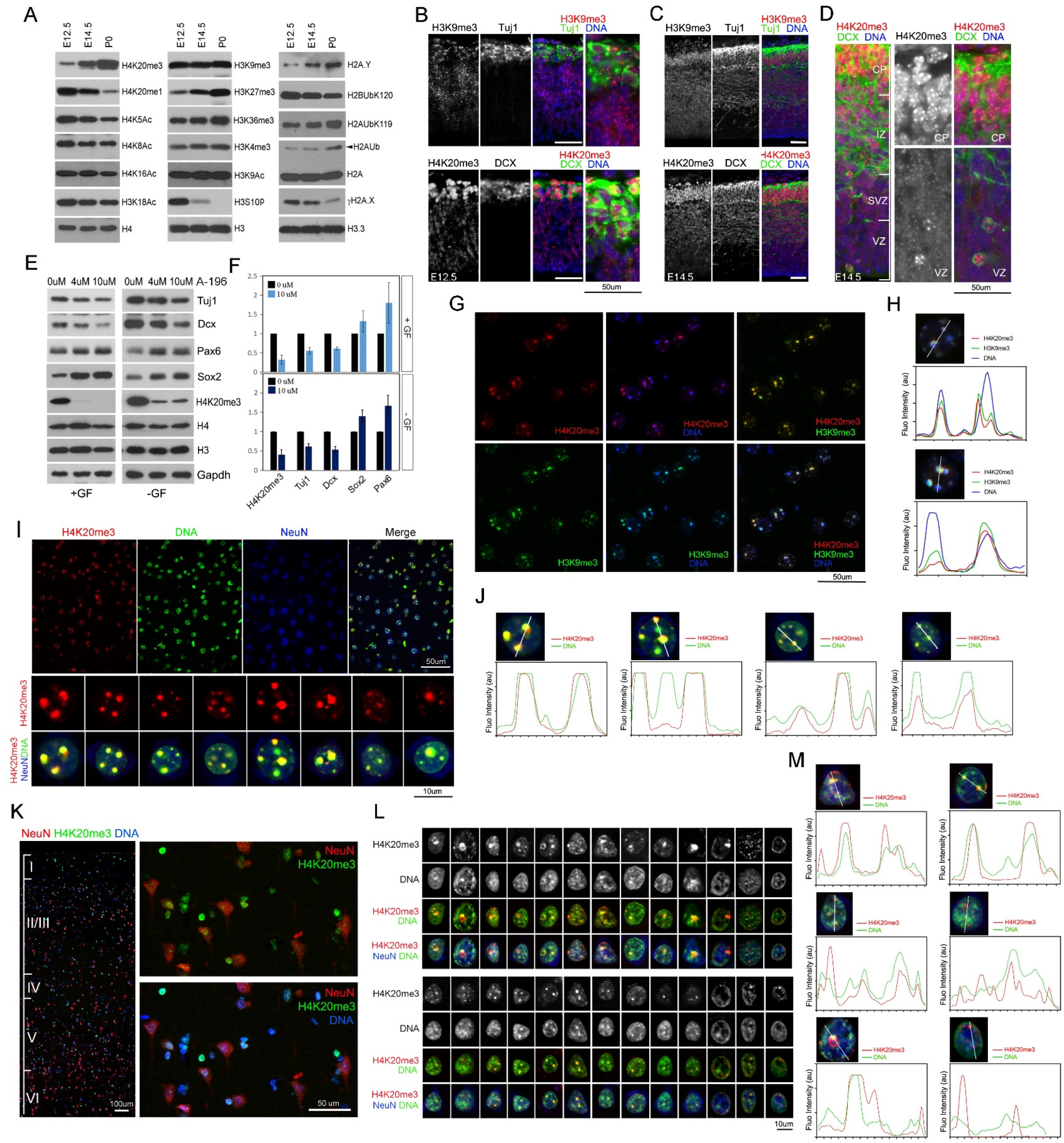
H4K20me3 modification of heterochromatin in neurogenesis yields diverse chromatin landscapes in neocortical neurons. (A) IB of histone extracts from developing murine cerebral cortical tissues at E12.5, E14.5, and P0. Note the progressive increase of H4K20me3, H3K27me3, and H2Aub during the course of cortical neurogenesis. (B-D) IH analysis of embryonic cerebral cortical sections with antibodies against H3K9me3 and H4K20me3. Sections are co-stained with new neuron markers Tuj1 or DCX to reveal the intense H4K20me3 foci in neurons of the nascent cortical plate at E12.5 (B), as well as in DCX+ newborn neurons in the VZ/SVZ, IZ and CP at E14.5 (C, D). Bars: 50 um. VZ: ventricular zone; SVZ: subventricular zone; IZ: intermediate zone; CP: cortical plate. (E, F) Representative IB images and quantifications of whole cell extracts from NPCs cultured in the presence or absence of growth factors (GFs). The cultures were treated with A-196 for 72 hours at the indicated concentrations. (G) Representative IH images of H4K20me3 (red)-H3K9me3 (green) double labeled neocortical sections of young adult mice. Nuclear DNA is stained by Hoechst 33342 (blue). (H) Line scan of H4K20me3, H3K9me3, and Hoechst 33342 fluorescence intensities in selected nuclei of mouse neocortical tissue, showing the co-localization of H4K20me3 with chromocenter and heterochromatin domains marked by H3K9me3. (I) Representative IH images of H4K20me3 (red)-NeuN (blue) double labeled neocortical sections of young adult mice. Nuclear DNA is stained by Hoechst 33342 (green). (J) Line scan of H4K20me3 and Hoechst 33342 fluorescence intensities in selected nuclear images of individual neocortical neurons. Note the diverse pattern of H4K20me3 distribution and occupancy with respect to chromocenters. (K) Representative IH images of NeuN (red)-H4K20me3 (green) double stained postmortem human cortical sections. Bars: 100um or 50um as indicated. (L) Selected nuclear images of individual neurons of postmortem human neocortical sections double stained by antibodies against H4K20me3 (red) and NeuN (blue). Nuclear DNA is stained by Hoechst 33342 (green). Note the greater heterogeneity of H4K20me3 landscapes and differential coverage of nuclear DNA by H4K20me3 in human than in murine brains. Bars: 10um. (M) Line scan of H4K20me3 and Hoechst 33342 fluorescence intensities in selected nuclear images of human postmortem neocortical neurons, showing hyper-heterogeneity of differential H4K20me3 occupancy with respect to chromocenters. See also Figure S1.

The remarkable increase in the intensity of H4K20me3 foci in newborn neurons prompted us to further ask whether H4K20me3 is required for neurogenesis. Tri- and di-methylation of lysine 20 in histone H4 is catalyzed by the histone methyltransferases Suv420h1 (KMT5B) and Suv420h2 (KMT5C) (Hahn et al., 2013; Schotta et al., 2008). We thus applied A-196, a potent and selective inhibitor of Suv420h1 and Suv420h2 (Bromberg et al., 2017), to NPC cultures and tested its effect on neuronal differentiation. We found that A-196 decreased the levels of new neuron makers Dcx and Tuj1 while increasing the levels of pro-NPC makers Pax6 and Sox2. This effect of A-196 on impeding neurogenesis was observed in both basal NPC cultures with low degrees of spontaneous neuronal differentiation and upon the induction of neuronal differentiation by growth factor withdrawal (Figure 1E, F). Therefore, the concomitant increase of heterochromatin compaction mediated by H4K20me3 is necessary for neurogenesis, through which post-mitotic neurons may gain stronger protection of repetitive sequences of the genome.

### H4K20me3 Marks Diverse Chromatin Landscapes in Neocortical Neurons

We further found that H4K20me3 not only compacts heterochromatin but also marks a wide array of chromatin landscapes of neocortical neurons. In these neurons, we observed that H4K20me3 foci largely overlapped with both the H3K9me3 condensates and the chromocenters containing A-T-rich satellite repeats that are stained intensively by the DNA dye Hoechst (Guenatri et al., 2004) (Figure 1G, H, Figure S1B). Furthermore, we found that, among neocortical neurons, H4K20me3-modified chromatin exhibited variable patterns and occupied deferent domains of the nucleus (Figure 1I, J). This suggests that H4K20me3 shapes diverse chromatin landscapes in various neurons besides reinforcing the compaction of the highly repetitive DNA elements to strengthen chromosome stability.

To further assess the association of H4K20me3 with the diversification of heterochromatin landscapes in neocortical neurons, we examined its distribution in human cerebral cortical tissue with the perception that the diversity of neocortical neurons increases substantially from mice to humans. Similar to what was observed in mice, H4K20me3-modified heterochromatin exhibited condensates of variable size throughout the nucleus of human cortical neurons (Figure 1K, L; Figure S1C). These condensates were enriched in the characteristic heterochromatin domains, showing preferential association with nuclear lamina and nucleoli (Figure 1L). However, the H4K20me3 landscape in human neurons is much more heterogeneous than in murine neurons. In human neurons, H4K20me3 condensates showed a wider size range, occupied various nuclear territories, and displayed differential coverage of the Hoechst-stained chromocenter DNA (Figure 1L, M). These variations in the abundance and distribution of H4K20me3 modified chromatin domains may be essential for increasing the epigenetic diversity, allowing an extraordinarily wide spectrum of chromatin landscapes to be established in neurons of the human cerebrum.

### Stalled DNA Replication in Nde1^-/-^ NPCs is Coupled to H4K20me3 Deficiency

Given the critical association of H4K20me3 with heterochromatin compaction in neocortical neurogenesis, we next determined whether defects in H4K20me3 underlie stalled heterochromatin replication that we previously observed in Nde1^-/-^ NPCs (Houlihan and Feng, 2014). DNA replication in S-phase starts with euchromatin and ends with heterochromatin. To determine where in S or G2 phase NPCs were, we labeled the NPCs progressing through S-phase by IdU-CldU sequential administrations to pregnant dams at E12.5 (Figure 2A). We then compared the number of H4K20me3-dense foci relative to the number of Hoechst-intense chromocenters in IdU and CldU labeled NPCs at early (IdU-;CldU+), mid (IdU+;CldU+), and late S-phase (IdU+; sparse CldU+ foci), respectively. Compared to WT NPCs, Nde1^-/-^ NPCs’ nuclei in S-phase exhibited an overall reduction in the intensity and number of H4K20me3 foci relative to the number of chromocenters (Figure 2B). By quantifying the numbers of H4K20me3 foci per chromocenter or per cell with respect to S-phase progression, we found that the numbers were comparable between WT and Nde1^-/-^ NPCs in early S-phase but were significantly reduced in Nde1^-/-^ NPCs in the mid and late S-phase or in those that showed stalled DNA replication (IdU+;CldU-, and localized in the S-phase zone)(Figure 2C-E). This result indicates that stalled heterochromatin DNA replication in Nde1^-/-^ NPCs was tightly coupled to compromised heterochromatin remodeling though H4K20me3, a necessary PTM for neuronal differentiation and genome protection.

**Figure 2.**
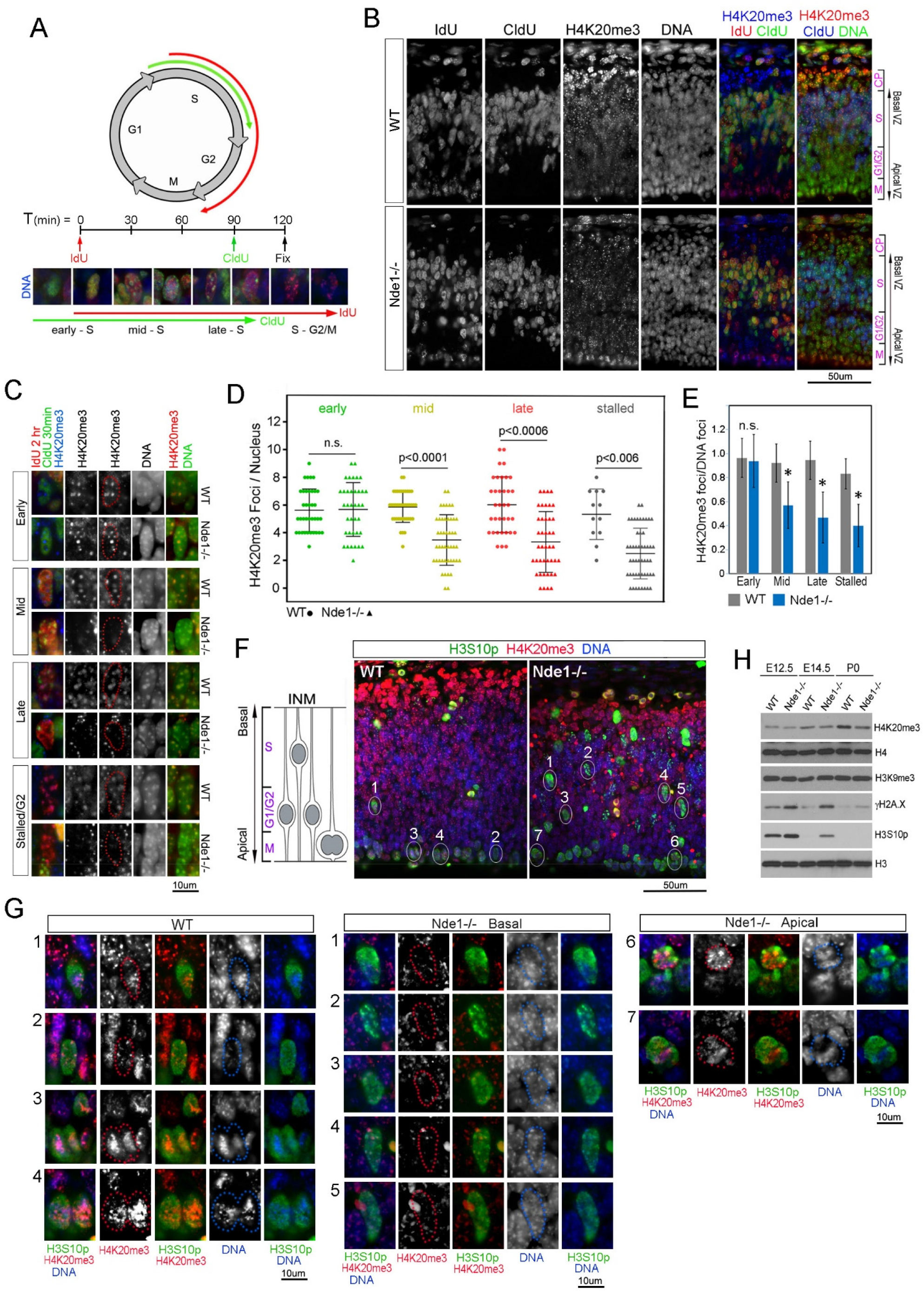
Stalled DNA replication, DSBs and G2/M arrests of Nde1^-/-^ NPCs are associated with compromised H4K20me3. (A) Diagram of IdU-CldU sequential labeling scheme for analyzing S-phase progression in embryonic cortical tissue. Pregnant dams were given IdU followed by CldU before embryos were fixed at E12.5. Embryonic NPCs were analyzed by IdU-CldU double IH staining. DNA were stained by Hoechst 33342. (B) Representative IH images of H4K20me3 in S-phase cortical NPCs of WT and Nde1^-/-^ embryos sequentially labeled by IdU and CldU as described in (A). DNA were stained by Hoechst 33342. VZ: ventricular zone; CP: cortical plate; S: S-phase zone; G2/M: G2 and M-phase zone. (C) Representative IH images of H4K20me3 in NPCs at early S-phase (IdU-;CldU+), mid S-phase (IdU+;CldU+), late S-phase (IdU+; sparse CldU+ foci), or stalled S-phase (IdU+; CldU- and in the S-phase zone of basal VZ as opposed to where G2/M nuclei localize at the apical VZ) at E12.5. DNA were stained by Hoechst 33342. H4K20me3 and Hoechst 33342 double stained images are included to reveal the H4K20me3 occupancy at chromocenters. Nuclear territory is marked by red dots on H4K20me3 images according to DNA stains by Hoechst 33342 (middle column). Note the weak and diffuse H4K20me3 immunosignals in late S-phase and DNA replication-stalled Nde1^-/-^ NPCs. (D) Numbers of H4K20me3+ foci per nucleus in NPCs of E12.5 embryos at early, mid, late, and stalled S-phase. P values by two-sample Kolmogorov-Smirnov tests are indicated. (E) Ratios of H4K20me3 to Hoechst 33342+ foci in nuclei of early, mid, and late S-phase or DNA replication stalled cortical NPCs at E12.5. Mean**±**SD. * p< 0.0001, by student’s t tests. (F) Representative IH images of E12.5 cortical sections that were doubled labeled by antibodies against phospho-Histone H3 Ser10 (H3S10p, in green) and H4K20me3 (in red). DNA was stained by Hoechst 33342 (in blue). Diagram to the left illustrates cell cycle-dependent interkinetic nuclear migration (INM) of cortical NPCs. (G) High-magnification IH images of H4K20me3 in selected H3S10p+ nuclei shown in (F) (indicated by circles and numbers). Note the reduced number and intensity of H4K20me3 foci in Nde1^-/-^ nuclei retained in basal ventricular zone. (H) IB of histone extracts from cortical tissue at E12.5, E14.5 and P0, showing that Nde1 LOF results in decreased H4K20me3 with increased DSBs (γH2A.X) and G2/M arrest (H3S10p). See also Figure S2.

A characteristic phenotype shown by Nde1^-/-^ NPCs was the mis-localization of mitotic nuclei and skewed mitotic spindles (Feng and Walsh, 2004). Given that constitutive heterochromatin encompasses primarily the centromere and pericentromere regions, we next sought to determine whether such mitotic defects of Nde1^-/-^ NPCs could arise from aberrant H4K20me3 modification in heterochromatin replication of S-phase. NPCs in WT embryos are known to undergo interkinetic nuclear migration (INM) and localize their nuclei at the apical end adjacent to the brain ventricles during mitosis (Takahashi et al., 1993), but mitotic nuclei of Nde1^-/-^ NPC frequently failed to localize at the apical end (Feng and Walsh, 2004). We, therefore, immuno-labeled mitotic NPCs by phospho-histone H3 Ser10 (H3S10p) and examined H4K20me3 in these cells. We found that H4K20me3 was largely condensed in H3S10p-labeled WT and those Nde1^-/-^ nuclei that were appropriately localized to the apical end. However, the H4K20me3 foci were substantially weakened, reduced in numbers, or absent in those H3S10p-labeled Nde1^-/-^ NPCs with basally mis-localized nuclei (Figure 2F, G). This strongly suggested that the mitotic defects in Nde1^-/-^ NPCs were coupled to compromised H4K20me3.

We confirmed that the overall level of H4K20me3 was reduced in the cortical tissue of Nde1^-/-^ embryos compared to that of their WT littermates by IB analyses of cerebral cortical histone extracts. Notably, decreased H4K20me3 abundance was inversely correlated with the level of histone PTMs associated with mitosis (H3S10p) and DNA double strand breaks (γH2A.X) in Nde1^-/-^ cortices (Figure 2H), supporting the notion that DSBs and mitotic arrests in Nde1^-/-^ NPCs result from aberrant heterochromatin remodeling and replication in neurogenic cell cycles. Taken together, our findings demonstrate that Nde1’s essential role in neurogenesis lies in ensuring effective epigenetic heterochromatin compaction, which is necessary for both patterning the chromatin landscape and for endowing genome stability of neocortical neurons. Considering the increasing demand on both epigenetic diversity and genome protection of neocortical neurons in mammalian evolution, this role of NDE1 should be more indispensable in humans than in mice.

### NDE1/Nde1 is an IDP that can Phase Separate and Bind to Constitutive Heterochromatin

To determine how NDE1 mediates heterochromatin remodeling and stability, we next investigated its molecular dynamics. At present, the mechanism that directs the de novo assembly of heterochromatin in vivo remains elusive, but recent work suggests that the formation of compact heterochromatin domains relies on the local accumulation of principle epigenetic heterochromatin writer molecules via a liquid-liquid phase separation (LLPS) (Larson et al., 2017; Strom et al., 2017). LLPS is generally believed to result from multivalent interactions between intrinsically disordered proteins (IDPs). This is well in line with the structure of NDE1, as both human and murine paralogs of NDE1/Nde1 contain an extended coiled-coil domain through which it can self-associate and/or mediate multiple molecular interactions in a concentration dependent manner. Predictor of natural disordered regions (PONDR) also indicates that over 90% of NDE1/Nde1 protein is disordered, containing large intrinsically disordered regions (IDRs) (Figure 3A). This molecular feature also fully agrees with our previous data showing that Nde1 is a dynamic scaffold protein that can partition into various subcellular domains including the nucleus (Houlihan and Feng, 2014; Pawlisz and Feng, 2011). We thereby tested if NDE1/Nde1 can phase separate by expressing GFP-tagged Nde1 and evaluating its dynamic subcellular localization and biophysical properties in forming a condensed liquid state. We observed that GFP-Nde1 displayed a variety of patterns with respect to subcellular distribution. At higher concentrations, a majority of GFP-Nde1 formed membrane-less condensates in both the cytoplasm and the nucleus (Figure 3B). These condensates exhibited liquid-like features, forming spherical droplets, showing rapid movement and exchange kinetics, and undergoing fusion to coalesce into larger spheres or immiscible compartments (Figure 3C-F, supplemental videos). We monitored GFP-Nde1 turnover within droplets by fluorescence recovery after photobleaching (FRAP) and observed a quick recovery of both cytoplasmic and nuclear GFP-Nde1 after bleaching (Figure 3G, H). These dynamic cell molecular and biophysical properties enable NDE1/Nde1 to nucleate phase-separated compartments, making it an excellent candidate to facilitate the molecular assemblies necessary for heterochromatin condensation.

**Figure 3.**
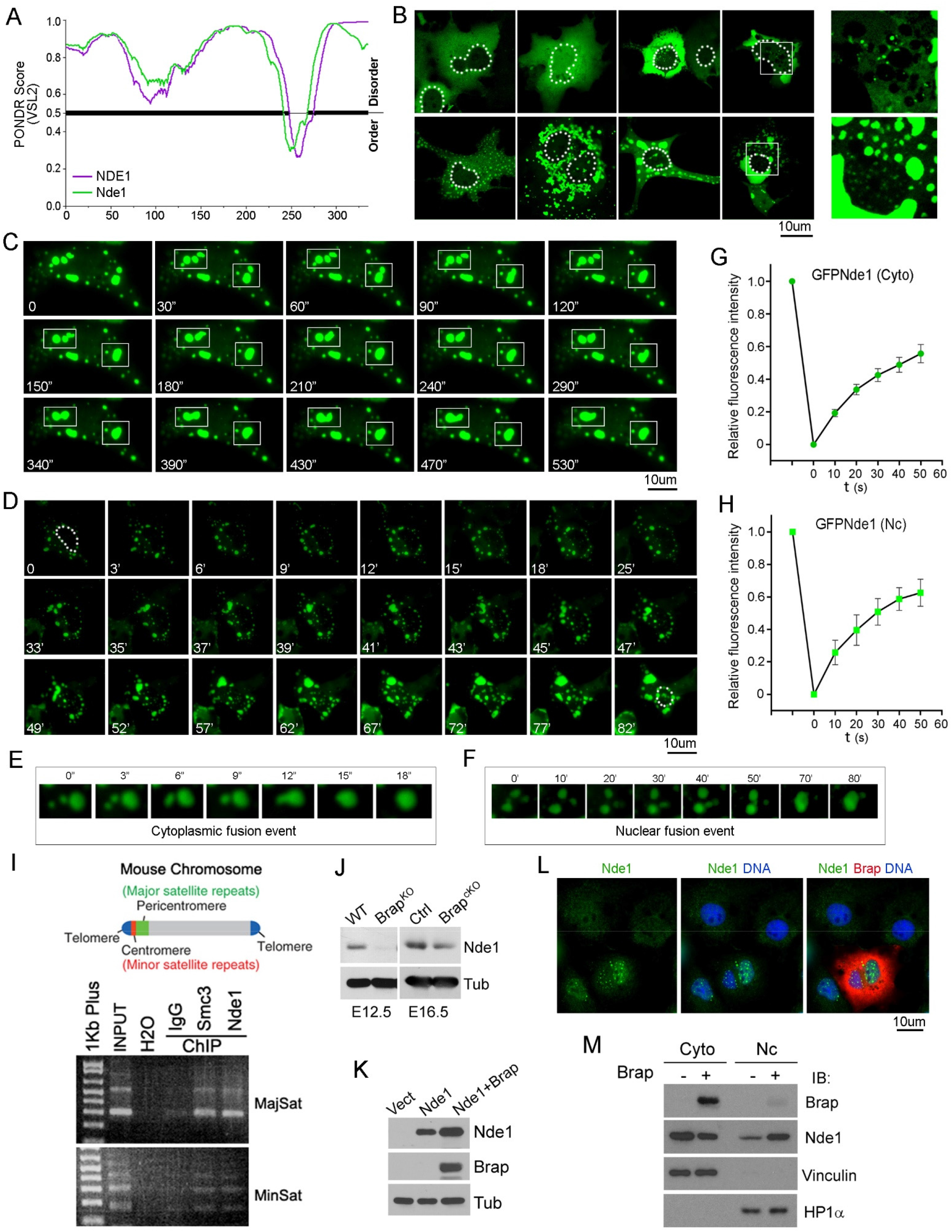
Nde1 is an IDP that can form spherical condensates through LLPS and directed to the nucleus by Brap. (A) Intrinsic disordered regions of human NDE1 and murine Nde1 proteins (by Predictor of Natural Disordered Regions). VSL2 scores are shown on the y axis, and amino acid positions are shown on the x axis. (B) Fluorescence confocal microscopy images of EGFP-Nde1 expressed in Cos7 cells, revealing the variety of distribution patterns. The position of the nucleus is marked by dotted lines, high-magnification images of the boxed areas are included. (C) Time-lapse images of Cos7 cells expressing EGFP-Nde1 (laser excitation every 10 seconds for the time indicated), showing dynamic fusion and redistribution. The position of the nucleus is marked on the panel of t=0. (D) Time-lapse images of Cos7 cells expressing EGFP-Nde1 (laser excitation every 10 minutes for the time indicated), showing the nuclear import and the formation of nuclear condensates. The position of the nucleus is marked on panels of t=0 and t=82’, respectively. (E) An example of droplet fusion event of cytoplasmic EGFP-Nde1. (F) An example of droplet fusion event of nuclear EGFP-Nde1. (G) FRAP of cytoplasmic GFP-Nde1 condensates. Shown is the average signal intensity (n=7) relative to the pre-bleaching signal (y axis) versus time relative to photo-bleaching (x axis). (H) FRAP of nuclear GFP-Nde1 condensates. Shown is average signal intensity (n=7) relative to the pre-bleaching signal (y axis) versus time relative to photo-bleaching (x axis). (I) A diagram of mouse chromosome structure and results of ChIP-PCR analyses of mouse embryonic E12.5 cortical tissue, indicating the physical association of Nde1 and Smc3 of the cohesin complex with MajSat and MinSat repeats of constitutive heterochromatin. (J) IB analysis of embryonic cerebral cortical tissue of Brap^KO^ (Brap^-/-^) and Brap^cKO^ mice and their WT or control littermates, showing that the level of Nde1 protein is decreased by Brap abrogation. This indicates the requirement of Brap for the stability of Nde1. (K) IB analysis of whole protein extracts from HEK293 cells transfected by recombinant Nde1 and Brap, revealing that the level of Nde1 is enhanced by Brap expression. (L) A representative IF image of Cos 7 cells transfected by recombinant Brap and double stained with antibodies against Nde1 (green) and Brap (red). Nuclear DNA is stained by Hoechst 33342 (blue). It demonstrates that the nuclear localization of endogenous Nde1 is enhanced in cells with Brap over-expression. (M) Cell fractionation and IB analyses of Cos7 cells transfected with recombinant Nde1 and Brap, showing increased Nde1 in the nuclear fraction of cells with Brap expression. See also Supplemental Videos.

We further determined the physical association of Nde1 with heterochromatin to assess Nde1’s role in driving heterochromatin remodeling. We have previously identified a direct interaction of Nde1 with the cohesin complex and showed that Nde1 LOF resembles the phenotypes of cohesinopathies (Houlihan and Feng, 2014). As the cohesin complex binds the pericentric heterochromatin with the highest affinity, we tested if Nde1 interacts with the centric- and pericentric-bond cohesin by chromatin IP (ChIP) of the embryonic cortical tissue. As expected, ChIP with antibodies against Nde1 and the Smc3 of cohesin each brought down pericentromeric major satellite and centromeric minor satellite repeats (Figure 3I). This robust interaction of Nde1 with satellite repeats suggests that Nde1 collaborates with cohesin to mediate the molecular assembly, compaction, and protection of centric and pericentric heterochromatin.

### The Nuclear Partition of NDE1/Nde1 is Directed by BRAP/Brap

The primary sequence of NDE1/Nde1 predicts a partial nuclear localization signal (NLS score of 5.8 by cNLS Mapper, and a score of 0.61 by NucPred), yet Nde1 was localized to the nucleus of NPCs during the S-phase of neuronal lineage restrictive cell cycles (Houlihan and Feng, 2014). As nuclear import would be required for Nde1 to form nuclear condensates and mediate heterochromatin compaction, we next sought to determine the mechanism by which NDE1/Nde1 is directed to the nucleus. One of Nde1’s essential molecular partner in central nervous system (CNS) development is BRAP/Brap, a BRCA1 associated protein capable of interacting with the nuclear localization signals (NLS) through which it regulates the cytoplasmic retentions vs. nuclear imports of its interacting molecules (Lanctot et al., 2013; Li et al., 1998). Nde1 and Brap have both common and differential molecular interactors. Brap has broader functions than Nde1 in regulating cell signaling and ubiquitin-mediated protein turnover (Lanctot et al., 2017; Lanctot et al., 2013), yet we found the level of Nde1 was reduced in the developing cortical tissue of mice with Brap knockout (Brap^-/-^) or Emx1-Cre-mediated Brap conditional knockout in dorsal telencephalic NPCs (Brap^flox/flox;Emx1Cre^, referred to as Brap^cKO^ hereafter) (Figure 3J). Conversely, we overexpressed recombinant Brap in cell cultures and found it was able to elevate not only Nde1’s overall level but also its nuclear partition (Figure 3K-M). These data suggested that while the stability of cytoplasmic Nde1 relies on its interaction with Brap, Brap can also promote Nde1’s nuclear partition. Given that Brap can undergo autoubiquitination to up- and down-regulate its own levels in response to growth signals (Lanctot et al., 2013; Matheny et al., 2004), the unique characteristics of Brap are likely both necessary and sufficient for stabilizing Nde1 and directing Nde1’s nuclear import in response to various physiological and pathological conditions.

To ascertain that Brap is essential for Nde1’s role in heterochromatin replication and remodeling, we determined whether Brap LOF phenocopies the heterochromatin defects of Nde1^-/-^ NPCs. We have previously shown that Brap^-/-^ NPCs have a shortened G1 phase but prolonged S phase in cell cycles towards neuronal differentiation, and that DNA replication in Brap^-/-^ NPCs stalls in late S phase similar to Nde1^-/-^ NPCs (Lanctot et al., 2017). To test whether the stalled DNA replication in Brap^-/-^ NPCs phenocopied Nde1^-/-^ NPCs with compromised H4K20me3, we analyzed the H4K20me3-dense foci and chromocenters in early, mid, late, and stalled S-phase NPCs identified by sequential IdU-CldU labeling (Figure S2A, B). As expected, we observed a significant decrease in the number of H4K20me3 foci in those Brap^-/-^ NPCs that were in late S-phase or showed stalled DNA replication at heterochromatin domains (Figure S2A-D). Furthermore, the overall level of H4K20me3 is reduced in nascent cortical neurons of Brap^-/-^ and Brap^cKO^ embryos (Figure S2A, E, F). This phenotype in heterochromatin replication and H4K20me3 modification closely resembles what we observed in Nde1^-/-^ NPCs. The phenocopy of Nde1 and Brap LOF strongly supports their functional collaboration in heterochromatin establishment during cortical neurogenesis.

### Heterochromatin and Nuclear Architecture Aberrations in Nde1 and Brap Deficient Neocortical Neurons

As the epigenetic modification of constitutive heterochromatin is expected to be stably maintained, we asked whether the failures in H4K20me3 patterning that we observed in neurogenic Nde1^-/-^ and Brap^cKO^ NPCs lead to chromatin defects of their daughter neurons. We first compared nuclei of newborn neurons in the cortical tissue of WT and Nde1^-/-^ embryos by transmission electron microscope (TEM). We found that the most prominent gross defect displayed in some Nde1^-/-^ neurons was an enlarged heterochromatin island in the center of the nuclei (Figure 4A). Heterochromatization typically drives the nuclear compartmentation by localizing euchromatin in the interior and heterochromatin at the periphery in conventional nuclei. By having a large heterochromatin condensate in the nuclear center but little or no heterochromatin in diverse nuclear domains and nuclear periphery, these Nde1^-/-^ nuclei appear “inverted” with respect to chromatin compartments. Therefore, compromised heterochromatin replication and remodeling in Nde1^-/-^ NPCs can result in cortical neurons with global nuclear architecture alteration.

**Figure 4.**
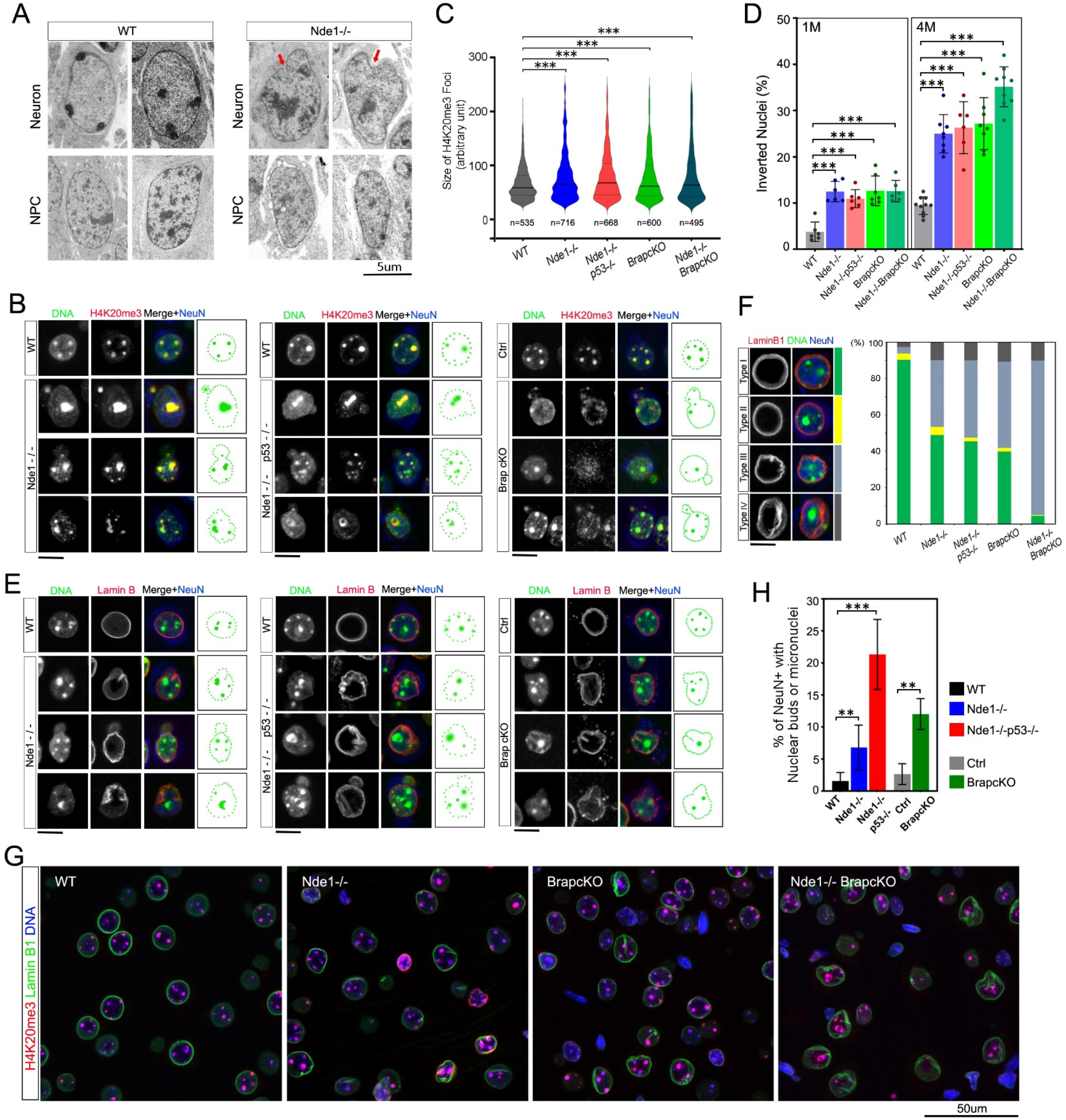
Nde1 and Brap LOF result in nuclear architecture aberrations in heterochromatin-associated domains. (A) Transmission electron microscopy (TEM) images of representative cortical NPCs and neurons in WT or Nde1^-/-^ embryos. Arrows indicate nuclear lamina aberrations. Bar: 5um. (B). IH images of individual nuclei from cortical sections of mice at weaning age (P22-28), showing diverse nuclear architecture and heterochromatin (marked by H4K20me3) aberrations in cortical neurons (NeuN+) of WT, Nde1^-/-^, Nde1^-/-^p53^-/-^, and Brap^cKO^ mice. Shown are representative images of nuclei from cortical sections of mice of indicated genotypes. n ≥ 5 for each genotype. (C) Violin plots of H4K20me3 foci size distribution on IH images of cortical neurons in mice of genotypes indicated at weaning age (P22-P28). The total number of foci quantified (by Image-Pro Premier) for each genotype (n ≥ 3 mice per genotype) are indicated. ***p<0.001 by two-sample Kolmogorov-Smirnov tests. (D) Quantification of inverted nuclei (%) in cortical tissues of mice of genotype indicated at weaning (1M) and 4 months (4M) of ages. Shown are Mean**±**SD and distributions of the percentage of inverted nuclei from different mice (n ≥ 3 mice per genotype). *** p<0.001 by student’s t-tests. (E) IH images of diverse nuclear architecture and nuclear lamina (marked by Lamin B1) in cortical neurons (NeuN+) of WT, Nde1^-/-^, Nde1^-/-^p53^-/-^, and Brap^cKO^ mice at weaning age (P22-28). Shown are representative images of nuclei from cortical sections of mice of indicated genotypes (n ≥ 5 mice for each genotype). (F) Quantification of nuclear inversion, nuclear lamina aberration, and the coincidence of nuclear inversion and nuclear lamina aberration in neocortical neurons of mice of indicated genotypes at 3-4 months of age (n ≥ 3 mice for each genotype). The percentages of four types of nuclei were scored. Type I: conventional nuclei with normal nuclear lamina; Type II: inverted nuclei with normal nuclear lamina; Type III: conventional nuclei with aberrant nuclear lamina; Type IV: inverted nuclei with aberrant nuclear lamina. (G) IH images of H4K20me3-Lamin B1 double stained neocortical sections of WT, Nde1^-/-^, Brap^cKO^, and Nde1^-/-^Brap^cKO^ mice of weaning age (P22-P28). Note the phenocopy and synergy of Nde1 and Brap LOF. Bar: 50um. H. Frequency of detectable nuclear buds or micronuclei in cortical neurons (NeuN+) of mice of genotypes indicated at weaning age (P22-P28). Mean**±**SD, **p<0.01, ***p<0.001 by student’s t-tests. Bars: 10um or as indicated. DNA was visualized by staining with Hoechst 33342. See also Figure S3.

We further performed immunohistological (IH) analyses to determine the effect of Nde1 LOF on the nuclear architecture of mature neocortical neurons at post weaning ages. We included mice with Nde1 and Trp53 double LOF (Nde1^-/-^p53^-/-^) in order to rule out the effect of loss of NPCs or neurons due to Trp53-dependent apoptosis (Houlihan and Feng, 2014). In addition, Brap^cKO^ mice along with their control littermates (Brap^flox/+ Cre-^ or Ctrl) were analyzed in parallel to test whether Nde1 and Brap act in the same pathway of heterochromatin regulation. A minimum of five mice of each genotype were analyzed using antibodies specific for heterochromatin and various nuclear landmarks. We first examined the size and distribution of heterochromatin condensates recognized by H4K20me3 immuno-signals. In most WT cortical neurons, H4K20me3 foci were relatively uniform in size and predominantly co-localized with the Hoechst-intense chromocenters (Figure 1E, Figure 4B, C). However, in Nde1^-/-^, Nde1^-/-^p53^-/-^, and Brap^cKO^ cortical neurons, H4K20me3 foci exhibited a more heterogeneous pattern, showing a broader range in size and subnuclear distribution, with some congregated in the nuclear center and some dispersed as speckles throughout the nucleus (Figure 4B, C). Coinciding with increased large H4K20me3 condensates, we observed a higher population of inverted nuclei in neocortical neurons of Nde1^-/-^, Nde1^-/-^p53^-/-^, and Brap^cKO^ mice than in WT mice (Figure 4D). While the fraction of inverted nuclei increased from weaning to adult ages in both WT and mutants, they remained significantly higher in mutants than in WT cortical tissues.

Inverted nuclei have been observed in retinal rod cells of nocturnal mammals and cells with lamin B receptor abrogation (Solovei et al., 2009; Solovei et al., 2013). They are believed to result from loss of heterochromatin attractions to the nuclear lamina. Indeed, inverted nuclei that we observed in newborn neurons of Nde1^-/-^ embryos displayed abnormal nuclear lamina, showing rumpled contour or invagination (Figure 4A arrows). Furthermore, we found inverted nuclei in mature neocortical neurons with Nde1 or Brap deficiency also coincided frequently with abnormal lamina revealed by lamin B1 immunoreactivities. Approximately half of the cortical neurons in Nde1^-/-^, Nde1^-/-^p53^-/-^, and Brap^cKO^ mice displayed aberrant nuclear lamina, showing lobules and blebs or invaginations and crossed pieces due to loss of structure integrity or misplaced lamina that bisects the nucleus (Figure 4E, F). We also generated mice with Nde1 and Brap double LOF in neocortical tissues (Nde1^-/-^Brap^cKO^) and found that nuclear lamina were abnormal in nearly all cortical cells of the double mutant mice (Figure 4G). Likewise, phenotypes in H4K20me3 heterogeneity and nuclear inversion were significantly stronger in Nde1^-/-^Brap^cKO^ double mutant mice than in mice with Nde1 and Brap single LOF (Figure 4D, F), which indicated the functional synergy of Nde1 and Bap in heterochromatin compaction and stability. Additionally, the nucleoli in neocortical neurons of Nde1^-/-^, Nde1^-/-^p53^-/-^, and Brap^cKO^ mice also displayed higher heterogeneity in size relative to that of WT mice (Figure S3A-D), further demonstrating that both Nde1 and Brap LOF can cause a global change in nuclear architecture that alters the chromatin landscape of neocortical neurons.

Consistent with a role of heterochromatin in maintaining genome stability, we detected nuclear buds and micronuclei in 10-20% of cortical neurons in Nde1^-/-^, Nde1^-/-^p53^-/-^, and Brap^cKO^ mice. This was significantly higher compared to the only 1-4% found in WT or Ctrl cortices (Figure 4 H). These gross nuclear anomalies are commonly associated with chromosomal fragmentation and mis-segregation. Therefore, our data together demonstrated that through proper heterochromatin establishment and/or maintenance, Nde1 and Brap are essential for protecting genome stability of neocortical neurons.

Despite profound nuclear architecture aberrance of neocortical neurons, hippocampal neurons were mildly affected by Nde1 or Brap LOF (Figure S3E-G). As the neocortex is evolutionarily more recent than the hippocampus, the stronger neocortical phenotype in neuronal heterochromatin aberrance is in alignment with the higher indispensability of NDE1 in generating evolutionarily advanced neurons with greater demands in heterochromatin diversity and stability.

### Nde1 LOF Results in Heterochromatin Instability and DSBs in Neocortical Neurons

Proper chromatin organization has been demonstrated to have a striking effect on mutation rates, with rates being significantly higher in heterochromatin than in euchromatin (Roberts and Gordenin, 2014; Schuster-Bockler and Lehner, 2012). Therefore, heterochromatin aberrations in Nde1^-/-^ and Brap^cKO^ neurons may be associated with heterochromatic DNA damage. To test this, we examined several markers for DSB in the cortical tissue of WT, Nde1^-/-^, Nde1^-/-^p53^-/-^, and Brap^cKO^ mice. We have shown previously that γH2A.X, a hallmark for DSBs, peaked in Nde1^-/-^ NPCs at the onset of neurogenesis but subsided in later embryonic and early postnatal periods (Houlihan and Feng, 2014). However, after the mutant mice aged to 3 months or older, we found γH2A.X became readily detectable again in the cortical tissue of Nde1^-/-^, Nde1^-/-^p53^-/-^, and Brap^cKO^ mice. Notably, γH2A.X immunosignals were found predominantly in neurons (NeuN+) but rarely in glia (NeuN-) (Figure 5A, B). Consistent with the presence of γH2A.X, 53BP1 and Brca1, the two essential molecules for DSB repair, were also upregulated in Nde1^-/-^ and Nde1^-/-^ p53^-/-^ compared to WT and p53^-/-^ cortical tissues (Figure 5C). Moreover, IB of cortical histone extracts not only confirmed the elevation of γH2A.X in the cortical tissue of Nde1^-/-^ and Nde1^-/-^ p53^-/-^ mice but also showed that the increased γH2A.X in Nde1^-/-^ and Nde1^-/-^p53^-/-^ mice were predominantly penta-ubiquitinated (Figure 5D, E). The dual PTM of phosphorylation and ubiquitination of histone H2A.X is a characteristic DNA damage response that we observed in Brap^cKO^ mice and postmortem brains of patients with Alzheimer’s disease(Guo et al., 2020). Therefore, heterochromatin aberrations resulting from Nde1 LOF can lead to genome instability and DSBs in adult cortical neurons with age-dependent progression and pathological impacts.

**Figure 5.**
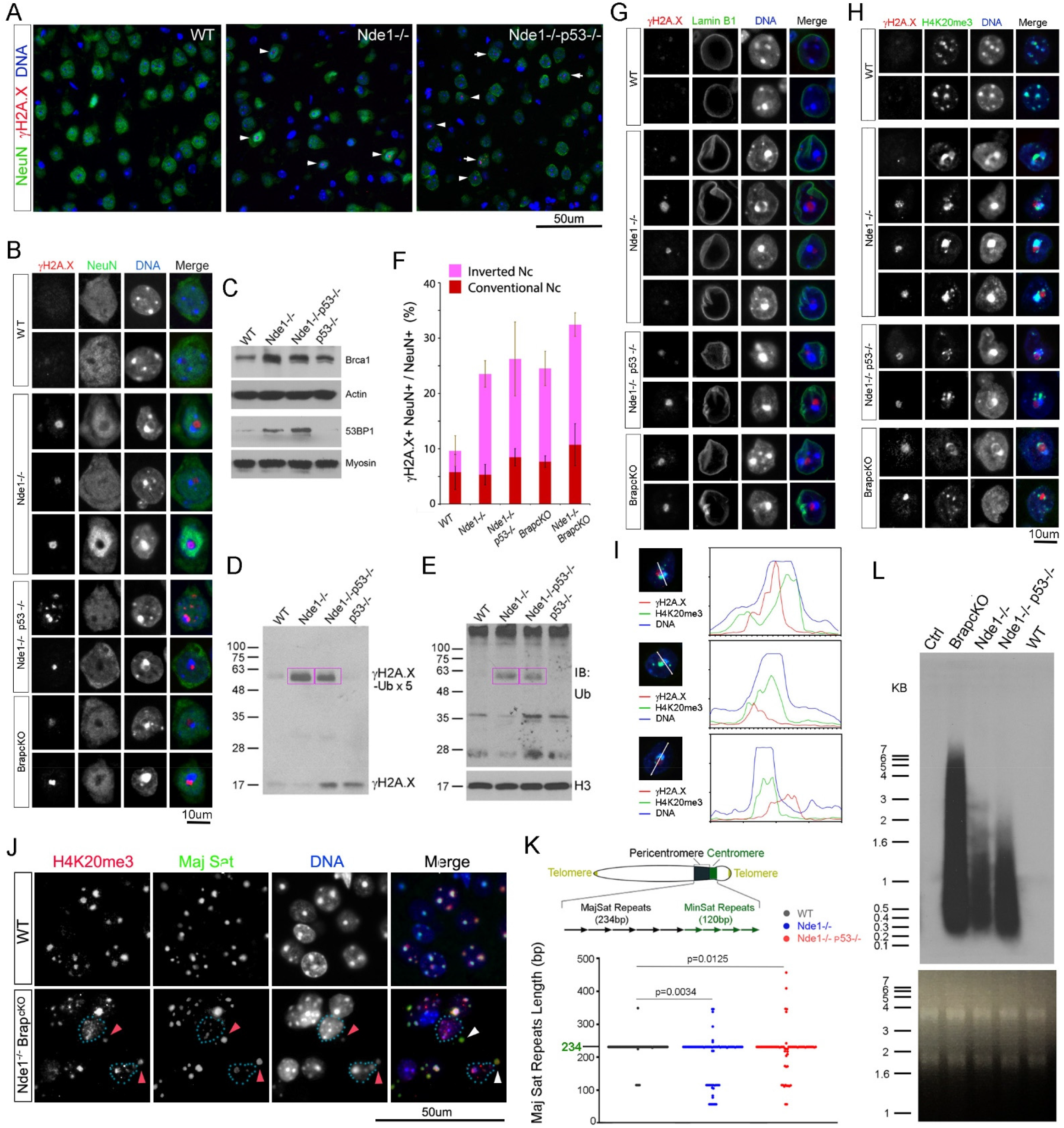
Nde1 and Brap LOF results in heterochromatin DSBs and increased transcription of pericentromeric satellite repeats. (A, B) Representative IH images of γH2A.X-NeuN double immuno-labeled neocortical sections of 4 month old WT, Nde1^-/-^, and Nde1^-/-^p53^-/-^ mice. Note that γH2A.X is shown specifically in the nucleus of NeuN+ cells (arrow heads). (C) IB of total cortical protein extracts from 4 month old mice of indicated genotypes. Note the substantial elevation of Brca1 and 53BP1 in Nde1^-/-^ and Nde1^-/-^p53^-/-^ cortical tissues. (D) IB of histone extracts with an anti-γH2A.X antibody, showing increased γH2A.X (∼17kDa) and penta-ubiquitinated γH2A.X (boxed, ∼60 kDa) in cortical tissues of Nde1^-/-^ and Nde1^-/-^p53^-/-^ mice of 4 months of age. (E) IB with an anti-ubiquitin antibody of the same blot shown in (D) after stripping off γH2A.X signals, confirming that the higher molecular weight band of ∼60 kDa recognized by anti-γH2A.X (boxed) contains ubiquitin modification. (F) Quantification (%) of neurons (NeuN+) showing γH2A.X and inverted nuclei in cortical tissue of 4 months old mice of genotypes indicated. Mean±SD, * p< 0.001 by student’s t tests. (G) Representative IH images of selected nuclei in γH2A.X-LaminB1 double immune-labeled neocortical neurons in adult (3-4 months old) WT, Nde1^-/-^, Nde1^-/-^p53^-/-^, and Brap^cKO^ mice. Note the coincidence of large γH2A.X foci in neurons with aberrant nuclear lamina. (H) Representative IH images of selected nuclei in γH2A.X-H4K20me3 double immuno-labeled neocortical neurons in adult (3-4 month old) WT, Nde1^-/-^, Nde1^-/-^p53^-/-^, and Brap^cKO^ mice. (I) Representative line scans of γH2A.X, H4K20me3, and Hoechst 33342 fluorescence intensities of selected individual nuclei, revealing the spatial proximity of γH2A.X with heterochromatin condensates. (J) Immuno-FISH analysis with a probe for major satellite DNA (green) and an antibody to H4K20me3 (red) on mouse neocortical sections. Shown are representative images, demonstrating the presence of pericentromeric major satellite repeats in the micronuclei of Nde1-Brap double deficient neocortical cells. Note the weaker H4K20me3 in the micronuclei (arrows). (K) PCR, cloning, and sequencing analyses of adult (2-4 months) cerebral cortical genomic DNA, showing increased polymorphic length variations of pericentromeric major satellite repeats in Nde1^-/-^, Nde1^-/-^p53^-/-^ cortices. Data were from n ≥ 3 mice per genotype. A total of 100 clones were sequenced from each genotype. P values by two-sample Kolmogorov-Smirnov tests are indicated. (L) Northern blot analysis of pericentromeric major satellite RNA in cerebral cortical tissues of 8-10 months old mice of indicated genotypes. The integrity and the amount of RNA loaded are shown by ethidium bromide stained gel below.

Further supporting a direct link between DSBs and abnormal chromatin structure in the mutant neurons, we found that more than 60% of cortical neurons that exhibited γH2A.X immunoreactivities also presented inverted nuclei (Figure 5F). In the cortical tissue of Nde1^-/-^, Nde1^-/-^p53^-/-^, and Brap^cKO^ mice, DSBs were prevalently detected in nuclei that exhibited severe nuclear lamina and heterochromatin aberrations (Figure 5G, H). In these nuclei, γH2A.X foci were found almost exclusively embedded in or flanking the heterochromatin condensates labeled by H4K20me3 and Hoechst-intense chromocenters (Figure 5H, I). This spatial proximity of DSBs with H4K20me3 condensates agrees with previous observations that DSBs are corralled to heterochromatin to increase protection against nuclease degradation of open chromatin ends and to facilitate homologous recombination (HR)-mediated repair (Ayoub et al., 2008; Ryu et al., 2015). Alternatively, this is also likely to indicate that DSBs occurred in heterochromatin DNA in mutant neurons, since the phosphorylation event resulting in γH2A.X can spread over hundreds of thousands to millions of bases (Gatti et al., 2012). To ascertain that the nuclear aberration indeed involves the physical damage of constitutive heterochromatin in Nde1^-/-^ and Brap^cKO^ neurons, we performed immunoFISH analysis using a probe for pericentromeric major satellite repeats (MajSat). We found that MajSat DNA was present in extranuclear micronuclei of Nde1^-/-^Brap^cKO^ double mutant cortical neurons, and that some micronuclei showed reduced H4K20me3 immunoreactivity (Figure 5J, arrows). Therefore, DSBs and chromatin fragmentations can occur in the satellite repeats of constitutive heterochromatin due to Nde1 and Brap deficiency. These results also suggested that compromised H4K20me3 led to gross chromosomal breaks and rearrangements.

To further test the impact of aberrant H4K20me3 and heterochromatin compaction on heterochromatin integrity, we asked whether Nde1 LOF results in changes in DNA sequence of heterochromatic repeats, since chromatin compaction by H4K20me3 prevents unscheduled homologous recombination of the underlying repetitive DNA (Bierhoff et al., 2014; Marion et al., 2009; Schotta et al., 2008; Wongtawan et al., 2011). In mice, the pericentromeric heterochromatin consists of large uninterrupted arrays of AT-rich major satellite repeats 234 base pairs in length that can extend more than 2 megabases per chromosome (Vissel and Choo, 1989). Since these extremely long repetitive DNA sequences are difficult to assess by Next Generation Sequencing, we used a PCR-based strategy to amplify MajSat monomers, cloned them, and examined their sequence integrity and polymorphism by Sanger sequencing. By analyzing PCR clones from genomic DNA isolated from cortical tissues of adult WT, Nde1^-/-^, and Nde1^-/-^p53^-/-^ mice at 2-4 months of age, we found 93% of the WT clones contained the standard monomers of MajSat repeats (77 of 83 clones from 4 mice), but only 70% of Nde1^-/-^ clones (58 of 83 clones from 4 mice) and 65.7% of Nde1^-/-^p53^-/-^ clones (46 of 70 clones from 3 mice) contained canonical MajSat monomers of 234 base pairs. The rest of the clones showed irregular length, which indicated that Nde1 LOF resulted in significant alterations in the sequence and structure of MajSat repeats (Figure 5K). These MajSat sequence variants support the notion that failures in heterochromatin compaction can lead to undesired recombination of repetitive DNA elements in Nde1 deficient cortical tissues.

### High MajSat Transcription from Heterochromatin Derepression in Nde1^-/-^ Cortical Tissue

Heterochromatin compaction is a key mechanism for repressing the transcription of its underlying repetitive DNA sequences to maintain their stability. The sequence variations of MajSat repeats along with the presence of MajSat DNA in micronuclei suggested compromised compaction and silencing of the MajSat repeats in Nde1^-/-^ cortices. We thus tested whether the heterochromatic DNA structure aberrance and DSBs in Nde1^-/-^ cortices were associated with the derepression of satellite transcription. By northern blotting analysis, we detected high levels of major satellite-derived transcripts in Nde1^-/-^, Nde1^-/-^p53^-/-^, and Brap^cKO^ cortical tissue (Figure 5L). These transcripts ranged from 200 to 6 kilobases, which was reminiscent of the aberrant overexpression of satellite repeats in mouse pancreatic ductal adenocarcinomas (Ting et al., 2011). As MajSat RNA transcripts were shown to be enriched in BRCA1 deficient breast cancers and acted as potent inducers of DNA damage (Zhu et al., 2018), the accumulation of pericentromeric satellite transcripts may account for DSBs and chromatin fragmentations in Nde1 and Brap deficient neurons.

### Progeria-like Histone PTMs, Neuroinflammation, and Neurodegeneration in Nde1^-/-^ Brains

The heterochromatin aberrations that we observed in the Nde1^-/-^ cortical tissue were reminiscent of the chromatin signatures of cancer and progeroid syndromes. However, there were no tumor-like phenotypes in the brain of Nde1^-/-^ and Nde1^-/-^p53^-/-^ mice, though some mutant mice that died between 3 and 12 months showed tumors outside of the CNS. This suggested that the consequence of nuclear architecture aberration and DSBs in postmitotic neurons is premature aging. This is supported by our study of Brap^cKO^ mice, which revealed a significant lifespan shortening along with accelerated cellular senescence and neurodegeneration (Guo et al., 2020). Therefore, we tested whether Nde1 LOF shared the characteristic histone PTMs of premature aging in progeroid syndromes. Hutchinson-Gilford Progeria syndrome (HGPS) is caused by *LMNA* mutations that affect lamin A processing. The aberrant lamin A proteins destabilize nuclear lamina and result in DSB accumulation similar to what we observed in Nde1 and Brap deficient cortical neurons (Liu et al., 2006; Scaffidi and Misteli, 2005). The nuclear and chromatin defects of HGPS are characterized by H4K20me3 upregulation (Arancio et al., 2014; Shumaker et al., 2005), whereas loss of H4K20me3 is a hallmark of cancer (Fraga et al., 2005; Yokoyama et al., 2014). Similarly, despite an initial H4K20me3 reduction in the developing cortices (Figure 2H, Figure S2F), Nde1^-/-^ and Brap^cKO^ mice showed higher levels of H4K20me3 in the adult and aging cortical tissue compared with their WT or control counterparts (Figure 6A). This age-dependent H4K20me3 upregulation in Nde1^-/-^ and Brap^cKO^ mice resembled the histone PTM changes of HGPS. It is also likely associated with cellular senescence (Nelson et al., 2016), and/or acts as a compensatory response to suppress the recombination and break of satellite repeats.

**Figure 6.**
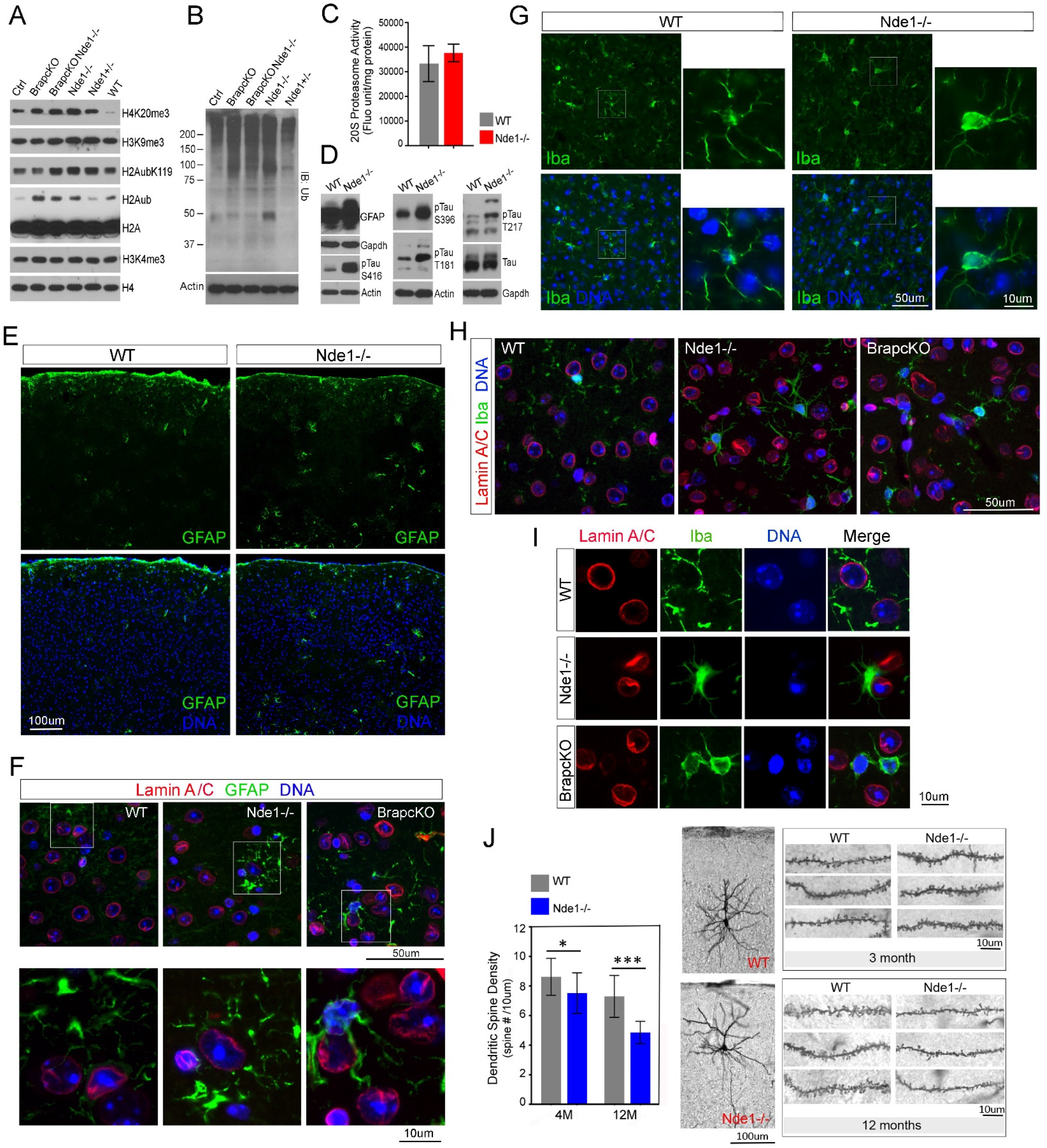
Nde1 LOF induces progeria and aging associated histone PTMs, neuroinflammation, and neurodegeneration. (A) IB of histone extracts from cortical tissues of mice of indicated genotypes at 4 months of age. Shown are representative images. Note the substantial increase in H4K20me3 and H2Aub in the cortical tissues with Nde1 or Brap LOF. (B) Ubiquitin IBs of cortical tissue total protein extracts from mice of indicated genotypes at 4 months of age. Note the accumulation of poly-ubiquitinated proteins in the cortical tissues with Nde1 or Brap LOF. (C) Proteasome activity analysis of WT and Nde1^-/-^ cortical tissues. Samples were prepared from 8-10 month old WT and Nde1^-/-^ mice. 20S proteasome activity was determined by assessing the amount of Suc-LLVY-AMC cleavage. (D) IB of cortical tissue total protein extracts from mice of indicated genotypes at 10 months of age. Shown are representative images. Note the increased levels of GFAP and phospho-tau in Nde1^-/-^ compared to WT mice. (E) IH images of GFAP (green) labeled cortical sections of WT and Nde1^-/-^ mice at 12 months of age. Shown are representative images. Note the increased and enlarged GFAP+ cells in Nde1^-/-^ mice compared to their WT littermates. Bar: 100um. (F) IH images of GFAP (green)-Lamin A/C (red) double immuno-labeled cortical sections of mice of indicated genotypes at 6 months of age. Shown are representative images. High-magnification views of the boxed areas are included. Note the spatial coincidence of aberrant nuclei and reactive astrocytes. Bars: 50um or 10um as indicated. (G) IH images of Iba (green) labeled cortical sections of WT and Nde1^-/-^ mice at 12 months of age. Shown are representative images. High-magnification views of the boxed areas are included. Note the morphological difference of microglia between WT and Nde1^-/-^ mice. Bars: 50um or 10um as indicated. (H) IH images of Iba (green)-Lamin A/C (red) double immuno-labeled mouse cortical sections of indicated genotypes at 6 months of age. Note the spatial coincidence of aberrant nuclei and activated microglia with amoeboid morphology. Bar: 50um. (I) High-magnification views of aberrant nuclei (marked by Lamin A/C, red) embraced by activated microglia (Iba, green). Bar: 10um. (J) Quantification of dendritic spine density (Mean ± SD) and representative images of Golgi-cox analyses of WT and Nde1^-/-^ mice at 3 and 12 months of age, showing age-dependent reduction in spine density in cortical layer 2/3 pyramidal neurons of Nde1^-/-^ mice. * p<0.01, *** p<0.0001 by student’s t tests. Bars: 100um or 10um as indicated. Nuclear DNA was stained with Hoechst 33342 in blue in all IH images.

The progeria-like H4K20me3 elevation in Nde1^-/-^ cortical tissue also coincided with increased histone H2A ubiquitination (H2Aub) (Figure 6A), a histone PTM that can occur on multiple sites of H2A in a context dependent manner to mediate transcriptional suppression and/or DNA repair (Uckelmann and Sixma, 2017). Ubiquitination on K119 by Polycomb repression complex 1 (PRC1), known to promote chromatin compaction and transcriptional silencing, was also reported to be a conserved biomarker for aging (Yang et al., 2019). Our data showed that histone H2Aub is a hallmark phenotype of Brap LOF and a pivotal mechanism to promote neurodegeneration by inducing histone degradation, proteasome overflow, and leading to a backlog of poly-ubiquitinated proteins in the cortical tissue (Guo et al., 2020). In the cortical tissue of Nde1^-/-^ mice, while a change in H2Aub was not evident during embryonic and neonatal development, H2Aub was significantly elevated along with penta-ubiquitinated γH2A.X at ages of 3-4 months or older (Figure 6A, 5E). As both H4K20me3 and H2Aub are histone PTMs that promote heterochromatin compaction, their spatiotemporal coincidence with elevated γH2A.X and nuclear architecture aberrations in aging Nde1^-/-^ and Brap^cKO^ neurons are likely to reflect structural changes of heterochromatin to cope with recurrent DSBs in the heterochromatic repetitive DNA. However, such histone PTMs may also induce secondary epigenetic and proteomic alterations, leading to premature brain aging as observed in Brap^cKO^ mice (Guo et al., 2020).

To corroborate that the heterochromatin defects in Nde1^-/-^ neurons can prime premature brain aging, we examined several phenotypes associated with neurodegeneration. Similar to what we found in Brap^cKO^ mice, the high histone H2Aub in the cortical tissue of Nde1^-/-^ mice coincided with a buildup of poly-ubiquitinated proteins at ages older than eight to ten months, even though the proteasome activates remained unaltered (Figure 6B, C). Also resembling what we observed in Brap^cKO^ mice, the proteomic aberrance in Nde1^-/-^ cortical tissues was accompanied by astrogliosis, microgliosis, and hyper-phosphorylation of the microtubule associated protein tau, which are all brain pathology hallmarks in the preclinical stage of neurodegenerative diseases (Figure 6D-I). We found that Glial Fibrillary Acidic Protein (GFAP), of which upregulation is a marker of reactive astrogliosis, was higher in the neocortices of Nde1^-/-^ mice compared to their WT littermates (Figure 6D-F). We also examined microglia, the resident immune cells of the brain. In WT cortical tissues, most microglia, identified by Iba, showed a ramified resting state appearance with many thin processes extending from a small cell body (Figure 6G). In contrast, many microglia in Nde1^-/-^ cortices exhibited an amoeboid shape with enlarged cell body and thicker processes (Figure 6G-I). These morphology characteristics indicated that microglia in Nde1^-/-^ cortical tissue were induced to enter a highly active state by pathological lesions. Notably, we found that the reactive astrocytes and microglia surrounded or closely contacted those neurons with severe nuclear lamina aberrations (Figure 6F, H, I). These suggest that astrogliosis and microgliosis in Nde1^-/-^ neocortical tissue were induced by the abnormal build-up of polyubiquitinated proteins and their associated toxic metabolites resulting from chromatin impairments and DNA damage responses.

To ascertain that the combined defects, including heterochromatin instability, the backlog of polyubiquitinated proteins, tau hyperphosphorylation, and neuroinflammation, together promoted neurodegeneration, we examined the dendritic spines of neocortical neurons in aged Nde1^-/-^ mice, since dendritic spine dysfunction is considered critical for the pathogenesis of neurodegenerative diseases and highly correlated with cognitive decline of Alzheimer’s disease. We performed Golgi-Cox staining to visualize the morphology and dendritic spines of neocortical layer II/III pyramidal excitatory neurons in WT and Nde1^-/-^ mice. While the gross morphology and dendritic branching pattern were not obviously affected by Nde1 LOF, the density of dendritic spines in Nde1^-/-^ neurons were significantly decreased compared to that in WT neurons (Figure 6J). This reduction in spine density was relatively mild at 4 months but became substantial as Nde1^-/-^ mice further aged to 12 months (Figure 6J). The layer II/III pyramidal excitatory neurons of the neocortex are essential for neuronal networking across various cortical regions and between the two cerebral hemispheres. Loss of dendritic spines in these neurons would impair their synaptic transmission and compromise higher-order brain functions. Given that the microcephaly phenotype of Nde1^-/-^ mice was due to failures in generating neurons of neocortical layer II/III (Feng and Walsh 2004), progressive functional decline of these neurons during adulthood would further drive cognitive decline and accelerate neurodegeneration.

## DISCUSSION

Neurons in the neocortex represent the largest repertoire of cell diversity and functional complexity across all tissues. These cells cease dividing at birth but function constantly with respect to electrical and metabolic activities throughout the organism’s lifespan. These distinctive features make it both an extraordinarily essential and a daunting task to ensure both the epigenome diversity and genome stability in neocortical neurons. What we learned through studying the function of NDE1/Nde1 illuminates a mechanism by which the chromatin diversity and stability in these cells are co-established. Our data suggest that Nde1 is an essential player for both neocortical neurogenesis and heterochromatin patterning by H4K20me3, a PTM that is not only abruptly elevated in neurogenesis but also diversely distributed throughout neuronal populations in the cerebral cortex. Nde1 LOF compromises H4K20me3 and results in intractable DSBs in neocortical neurons. Of special interest is the pericentromeric MajSat repetitive elements regulated by Nde1. These satellite repeats are modified by H4K20me3, assemble into highly condensed constitutive heterochromatin, and remain permanently condensed during interphase of most somatic cells. This is extremely essential to ensure the structural integrity and transcriptional silencing of these highly repetitive yet important genomic regions. It is known that the pericentric repetitive sequences flank the centromeres and are bound by cohesin, playing a major role in maintaining sister chromatid cohesion around centromeres to prevent premature chromatid separation during mitosis (Bernard, Science 2001). We now show that these elements are protected additionally by the cohesin binding protein Nde1. Concomitantly, Nde1 also controls the epigenetic silencing and condensation of these repetitive elements to stabilize the heterogeneous chromatin landscapes in post-mitotic neocortical neurons. Therefore, the indispensable role of NDE1/Nde1 in neocortical development is to ensure the epigenetic remodeling and stability of pericentromeric constitutive heterochromatin and its underlying satellite repeats.

Satellite DNA sequences are the most dynamic element of genome evolution, presenting specific sequences and distribution patterns that are associated with polygenesis and species-identities (Thakur et al., 2021). However, the role of satellite repeats and their associated constitutive heterochromatin remains unclear, as their high repetitiveness makes them technically difficult to assess. Although the importance of these regions is becoming more recognized as of late as technology improves (Nurk et al., 2021), a lack of in vivo models to determine their functional impacts also hinders their study. Our finding of hyper-variability of heterochromatin landscape in human neocortical neurons as well as their physical and functional regulation by NDE1/Nde1 illuminates the possible role of satellite repeats in evolutionarily-mediated control of neocortical neuronal identity, diversity, and genome stability. Therefore, the increased chromatin complexity and epigenetic diversity impose a stronger demand on NDE1, making it more indispensable in humans than in mice.

Our finding provides critical insight into a potential mechanism underlying the common etiology of neurological and psychiatric disorders. Neuropsychiatric illnesses, such as schizophrenia, depression, autism, ADHD, and epilepsy, are diagnosed based on their clinical presentations as opposed to standardized criteria or objective parameters. However, with respect to clinical manifestation, each of these disorders presents highly heterogeneous symptoms. Conversely, many of these disorders share common symptoms and/or often show comorbidity with each other. Moreover, the clinical symptoms of neuropsychiatric disorders are also observed in neurodegeneration. As a matter of fact, patients with neurodegenerative disorders not only exhibit social behavioral deficits, but the neuropsychiatric symptoms are even considered relevant indexes for determination of the severity and progression of neurodegeneration. Although the complex traits of neuropsychiatric and neurodegenerative disorders are believed to result from multiple genetic determinants and their interaction with environmental factors, mutations of genes controlling the genome stability and epigenetic regulation can result in somatic chromatin aberrations and/or DSBs in various neurons and brain regions. Given the stochastic nature of somatic genetic and epigenetic alterations, heterogeneous clinical phenotypes are expected. This explains how aberrations in a single gene, *NDE1*, can give rise to multiple brain disorders with a wide spectrum of clinical symptoms but without necessarily reducing the number of neocortical neurons. Additionally, as NDE1 is a scaffold IDP that enables it to undergo multivalent interactions through LLPS, it can form biomolecular condensates encompassing numerous distinctive types of proteins and genomic regions. Therefore, either an increase or decrease in the abundance of NDE1 is expected to alter NDE1-dependent molecular condensates, causing substantial change in diverse cellular activities. This is likely the mechanistic basis of clinical observations that copy number variations caused by both microdeletion and microduplication of *NDE1* in 16p13.11 cause overlapping, yet diverse, neuropsychiatric syndromes.

## STAR METHODS

### KEY RESOURCE TABLES

Antibodies Table.

### EXPERIMENTAL MODEL AND SUBJECT DETAILS

#### Mice

Nde1 knockout (Nde1^-/-^), Brap knockout (Brap^-/-^), Brap floxed (Brap^flox/flox^), Trp53 knockout (p53^-/-^), and Emx1^cre^ mediated Brap conditional knockout (Brap^cKO^) mice have been described previously (Feng and Walsh, 2004; Houlihan and Feng, 2014; Lanctot et al., 2017). The Trp53 floxed mice (p53^flox/flox^) were purchased from JaxMice (stock # 008462) and used to generate Nde1^-/-^p53^cKO^ mice by crossing with Nde1^+/-^ and Emx1^cre^ mice (JaxMice stock # 005628). Nde1^-/-^Brap^cKO^ mice were generated by crossing Nde1^+/-^ with Brap^cKO^ mice. All mice used for this study were housed and bred according to the guidelines approved by the IACUC committees of Northwestern University and Uniformed Services University of Health Services in compliance with the AAALAC’s guidelines. Experiments were performed using littermates or age and genetic background matched control and mutant groups in both sexes. For timed matings, the day of vaginal plug was considered E0.5.

#### Human Cerebral Cortical Tissue Samples

Paraffin sections of human postmortem cortical tissue were obtained from the NIH NeuroBioBank at the University of Miami and were analyzed in compliance with NeuroBioBank ethical guidelines. Three neurologically normal individuals (HCTTA, HCTRY, and HCTPC) who died between the ages of 29 and 33 of cardiovascular causes were analyzed.

### EXPERIMENTAL MODEL AND SUBJECT DETAILS

#### Cell culture, Transfection, and Fluorescence Microscopy

Neural stem/progenitor cells were isolated from embryonic cortices at E12.5. Single cells were prepared and cultured in DMEM/F12 with N2 and B27 supplements, Penicillin-Streptomycin, Glutamine, Heparin, and growth factors (20 ng/ml EGF and 10 ng/mL FGF) at 37°C in 5% CO2. Cultures were inspected daily and split 1:3 to 1:5 every 3 days depending on cell density and speed of growth. The A-196 inhibitor was dissolved in DMSO and added to the culture medium at a final concentration of 4 uM or 10 uM for 3 days. Then cells were collected for total protein extraction and IB analyses. Control cells were treated with DMSO alone.

Cos7 cells from ATCC were cultured in DMEM supplemented with 10% fetal bovine serum. cDNA transfection was performed using Lipofectamine 2000 (Invitrogen) following manufacturer’s instructions. cDNA plasmids of Nde1, GFP-Nde1, and Brap were described previously (Feng et al., 2000; Lanctot et al., 2017). Immunofluorescence cell staining was carried out as described (Lanctot et al., 2017; Lanctot et al., 2013). Briefly, cells were fixed with either 4% formaldehyde or cold methanol, blocked in 1% BSA and 5mg/ml lysine, and immuno-stained in a buffer containing 25 mM Hepes, pH 7.4, 250 mM Sucrose, 25 mM KCl, 25 mM Mg(CH_3_COO)_2_, 1% BSA, and 0.25% Saponin.

#### Time-lapse Fluorescence Microscopy

For time-lapse imaging of GFP-Nde1, transfected cells grown in phenol red-free DMEM and 10% FBS on coverslips were assembled into a Pecon POC-R2 chamber (PeCon GmbH, Erbach, Germany). This chamber was maintained at 37L and 5% CO_2_ for the duration of the experiment. The chamber was placed on a Leica DMI 6000B inverted microscope controlled by Leica LAS X software (Leica Microsystems, Inc., Buffalo Grove, IL). Images of selected cells were recorded at 40x magnification using both transmitted light (using differential interference contrast [DIC]) and fluorescent light (through a filter cube appropriate for GFP) either every 10 seconds for a duration of 15 minutes or every 10 minutes for a period of 16 hours.

#### Fluorescence Recovery after Photo-bleaching (FRAP)

For FRAP experiments, GFP-Nde1 transfected cells grown in phenol red-free DMEM and 10% FBS on coverslips were assembled into a Pecon POC-R2 chamber (PeCon GmbH, Erbach, Germany). This chamber was maintained at 37L and 5% CO_2_ for the duration of the experiment. The chamber was placed on a Zeiss 700 confocal microscope (Zeiss Microscopy, Jena, Germany) and images were recorded using Zeiss Zen software using settings appropriate for GFP. Selected cells were imaged at 10x magnification. A small region in either the cytoplasm or nucleus was then bleached using the bleaching function of the software and subsequent images were recorded at intervals ranging from 5 to 15 seconds. For intensity analysis of bleached regions, image sequences were imported into FIJI (Schindelin et al., 2012). Regions of interest (ROIs) were defined in FIJI with care so that even if the area to be measured moved over the duration of the experiment, the measured area was always completely filled with fluorescence. Numerical measures of pixel intensity were then imported into Microsoft Excel for analysis (Microsoft, Inc., Redmond, WA)

#### Immunoblotting

Immunoblotting of total cell or tissue proteins was performed by extracting with boiling 2XSDS PAGE sample buffer (62.5 mM Tris-HCl, pH 6.8, 2.5% SDS, 0.7135 M β-mercaptoethanol, 10% glycerol, 0.002% Bromophenol Blue) to fully dissolve the tissue proteins, heating at 95°C for 10 min to ensure protein denaturation, and passing through syringes with a 29^1/2^ gauge needle three times to sheer nuclear DNA and obtain homogenous extracts. 10-30 ug of proteins were loaded on the gel for immunoblotting analysis. Loadings were adjusted and normalized by the total protein content according to Coomassie blue stain of the gel after SDS PAGE and by the level of housekeeping proteins.

#### Histone Extractions

Cell or tissues were re-suspended or homogenized and lysed in PBS containing 0.5% Triton X 100, 25 ug/ml leupeptin, 10 ug/ml Pepstatin A, 5 ug/ml Aprotinin, 10 mM Benzamidine, 2 mM PMSF, 10mM *N*-Ethylmaleimide (NEM), 10 mM iodoacetamide (IAA), and 0.02% NaN_3_ at a density of ∼10^7^ cells/ml. Nuclei were first collected by centrifuge at 2500 x g for 10 min at 4°C, washed once, and re-suspended in 0.2 N HCl at a density of 4×10^7^ nuclei per ml to extract histones overnight at 4°C. After clearing debris by centrifugation at 10,000 x g for 10 min at 4°C, the histone containing supernatants were neutralized with 2 M Tris Base. Protein concentration was determined by measuring absorbance at 280 nm. Extracted histones were stored in aliquots at −20°C.

#### Immunohistological Analyses

Immunofluorescence staining of brain tissue sections was carried out as described (Feng and Walsh, 2004). Briefly, mouse brains were fixed by transcardial perfusion with PBS and 4% paraformaldehyde and then processed in 12 um cryosections or 5 um paraffin sections. After treating with antigen unmasking solutions (Vector Labs), brain sections were blocked with 5% goat serum and incubated with primary antibodies in PBS, 0.1% Triton X100, and 5% goat serum at 4°C overnight, and followed by staining with fluorescence conjugated secondary antibodies and Hoechst 33342. Epifluorescence images were acquired with a Leica CTR 5500 fluorescence, DIC, and phase contrast microscope equipped with the Q Imaging Regita 2000R digital camera. Images were imported to Adobe Photoshop and adjusted for brightness and black values.

#### Ultra-Structural Analysis

Mouse embryos were fixed in 2% glutaraldehyde and processed for standard transmission electron microscopy analysis as described (Pawlisz and Feng, 2011). Specimens were examined with a JEOL 1220 transmission electron microscope equipped with Kodak digital camera. Cell type identities were determined based on the location of the cells, in which NPCs were in the ventricular zone and neurons were in the cortical plate.

#### Golgi-Cox Staining and Dendritic Spine Analysis

Mice were euthanized with CO_2_; brains were quickly dissected, rinsed with deionized water, immersed in impregnation solution, and processed using FD Rapid GolgiStain kit (FD NeuroTechnologies) according to manufacturer’s instructions. Stained sections were examined under a Leica DM5000 light microscope. Pyramidal neurons in the cerebral cortex and hippocampus regions were imaged with 40x objective and photographed. For dendritic spine density analysis, 16-20 pyramidal neurons in neocortical layer II/III of each mouse were randomly selected for assessment. The number of spines per 10 micrometers in secondary apical dendrites (branched from primary dendrites arising from the soma) was scored using the NIH Image J software.

#### Nucleotide Administration and S-phase Analysis

S-phase analysis was performed as described (Houlihan and Feng, 2014). Briefly, pregnant mice were injected intraperitoneally with 50 mg/kg of thymidine analogue IdU followed by thymidine analogue CldU 90 min after IdU injection. Then mice were euthanized 30 min after CldU injection for embryo extraction and histological preparation. Early S-phase cells were identified by CldU+ IdU-labeling. Mid S-phase cells were identified by CldU and IdU double labeling. Cells in late S-phase and G2-phase were identified by IdU+ CldU-labeling.

#### Chromatin Immunoprecipitation

Cerebral cortical tissue was dissected from wild type embryos at E12.5, fixed for 30 minutes in 2 mM disuccinimidyl glutarate and then for 10 minutes in 1% formaldehyde. Fixed tissue was suspended in 5 mM NaCl, 0.05 mM EDTA pH 7.5, 5 mM, 0.005% NP40, 0.01% Triton X-100 to isolate nuclei. Isolated nuclei were sonicated using a Diagenode Bioruptor. Sheared chromatin was incubated with antibodies against Nde1 and Smc3, respectively, then precipitated with M280 paramagnetic beads (ThermoFisher) coated with anti-rabbit IgG. DNA was isolated using MinElute PCR purification columns (QIAGEN). Primer sequences for ChIP-PCR are: Major Satellite: GACGACTTGAAAAATGACGAAATC and CATATTCCAGGTCCTTCAGTGTGC; Minor satellite: GAACATATTAGATGAGTGAGTTAC and GTTCTACAAATCCCGTTTCCAAC.

#### Northern Blot Analysis

RNA was extracted from mouse cortical tissue using TRIzol reagent (Thermo Fisher) according to the manufacturer’s protocol. After removing genomic DNA with TURBO DNase (Ambion), 2.5 ug of total RNA was mixed with formaldehyde and formamide denaturation buffer, incubated at 55°C for 15 min, and electrophoresed in a 1.2% agarose gel with 0.6M formaldehyde. RNA was transferred onto Amersham Hybond N+ membrane (GE) and crosslinked with ultraviolet light. The membrane was prehybridized in NorthernMax Hybridization Buffer (Ambion). Northern blot probe for major mouse satellite was generated by PCR of mouse genomic DNA with primers ATATTTCACGTCCTAAAGTGTG and GGCGAGGAAAACTGAAAAAG using the PCR DIG Probe Synthesis Kit (Roche). 0.1 nM probe was applied to the prehybridized membrane and hybridized in NorthernMax hybridization buffer (Ambion) at 42°C overnight. The membrane was washed twice with 2X SSC and 0.1% SDS for 15 min each at room temperature followed by washing with 0.2X SSC and 0.1% SDS at 65°C twice for 15 min each. Chemiluminescent detection of DIG signals on the membrane was performed using CSPD, ready-to-use (Roche) according to manufacturer’s instructions.

#### Mouse Major Satellite Genomic DNA Analysis

Genomic DNA was isolated from mouse cortical tissue using genomic DNA extraction kit (QIAGEN). PCR amplifications of major satellite repeats were performed using primers ATATTTCACGTCCTAAAGTGTG and GGCGAGGAAAACTGAAAAAG. The PCR products were cloned with the TOPO TA Cloning Dual Promoter Kit (Invitrogen). The resulting PCR clones were sequenced by Sanger sequencing using T7 and M13 reverse primers. 100 clones per genotype were randomly selected for sequencing analysis. All clones containing major satellite sequences were included in the analysis.

#### ImmunoFISH Analysis

Mouse brain sections (5um) on glass slides were de-paraffined and incubated in 10mM citric acid buffer at 95C for 30min. The slides were washed three times with PBS, treated with 0.1% Triton X100 in PBS for 15min and followed by two 2XSSC washes, a cold acetone treatment for 5 min, and three more 2X SSC washes for 5 min each at room temperature. Then the slides were treated or stored in 2XSSC and 50% formamide for longer than 4 hours.

The FISH probe for mouse major satellite DNA was amplified from the plasmid clone pγSat that contains eight copies of the 234bp mouse major satellite (γ-satellite) repeats using PCR Fluorescein Labeling Mix (Roche). Oligo primers used for PCR amplification are GACGACTTGAAAAATGACGAAATC, and CATATTCCAGGTCCTTCAGTGTGC. Pretreated tissue slides were transferred to hybridization buffer containing 50% formamide, 20mM sodium phosphate pH7.0 and 2xSSC, co-denatured with 100ng of DNA probe per slide at 80°C for 5 minutes, followed by hybridization at 37°C for 48–72 hours. After hybridization, slides were washed three time with 2XSSC at 37°C for 5 min, three times with 0.1XSSC and 0.1% NP-40 at 60°C for 5 min, and once with 2XSSC at room temperature for 5 min. Then the slides were transferred to PBS and processed for immuno-histological stains.

#### 20S Proteasome Assay

Proteasomes were extracted from mouse cortical tissue in a lysis buffer containing 50 mM HEPES, pH 7.5, 5 mM EDTA, 150 mM NaCl and 1% Triton X-100, 1 uM DTT and 2 mM ATP. Proteasome activities were determined using a 20S proteasome activity assay kit (Millipore APT280) according to manufacturer’s instructions. The amount of cleaved AMC fragment of Suc-LLVY-AMC was quantified using a CLARIOstar Plus plate reader (BMG LABTECH) at excitation (EX) = 380/emission (EM) = 460. Reaction mixtures were incubated with 10 μM lactacystin or MG132 before addition of fluorogenic substrates to ensure the specificity of the assays.

#### Quantification and Statistical Analysis

No statistical methods were used to predetermine sample size, while all experiments were performed with a minimum of three biological replicates and all cell counts were obtained from at least ten random fields. The experiments were not randomized; the investigators were not blinded to the sample allocation and data acquisition during experiments but were blinded in quantitative image analyses using the Image-Pro Premier (Media Cybergenetics) and the NIH Image J software.

All statistical analyses were done using GraphPad Prism 9.0 software. Data were analyzed by unpaired two-tailed Student’s t tests or by Kolmogorov-Smirnov tests for comparing differences between different genotypes. Differences were considered significant with a p value < 0.05.

## Supporting information

Antibody list

## ACKNOWLEDGEMENT

The authors wish to thank Sui Huang of Northwestern University for providing antibodies and for discussion. This work was supported by R01HD56380 and startup funds of Northwestern University and Uniform Services University of Health and Sciences to Y.F.

## AUTHOR CONTRIBUTIONS

Y. F conceptualized the project, designed and performed the experiments, interpreted the results, and wrote the manuscript. A.A.C designed and performed the experiments, interpreted the results, and wrote the manuscript; C.C.L performed experiments; Y.G. performed experiments;D.M. performed cell imaging and FRAP analyses; H.P performed experiments; Q.Z. performed experiments; X.Z assisted with experiments; M.L.D assisted with experiments.

## DISCLAIMER

The opinions, interpretations, conclusions and recommendation are those of the authors and are not necessarily endorsed by the U.S. Army, Department of Defense, the U.S. Government or the Uniformed Services University of the Health Sciences. The use of trade names does not constitute an official endorsement or approval of the use of reagents or commercial hardware or software. This document may not be cited for purposes of advertisement.

