## Supplementary Figures for "Nde1 is Required for Heterochromatin Compaction and Stability in Neocortical Neurons"

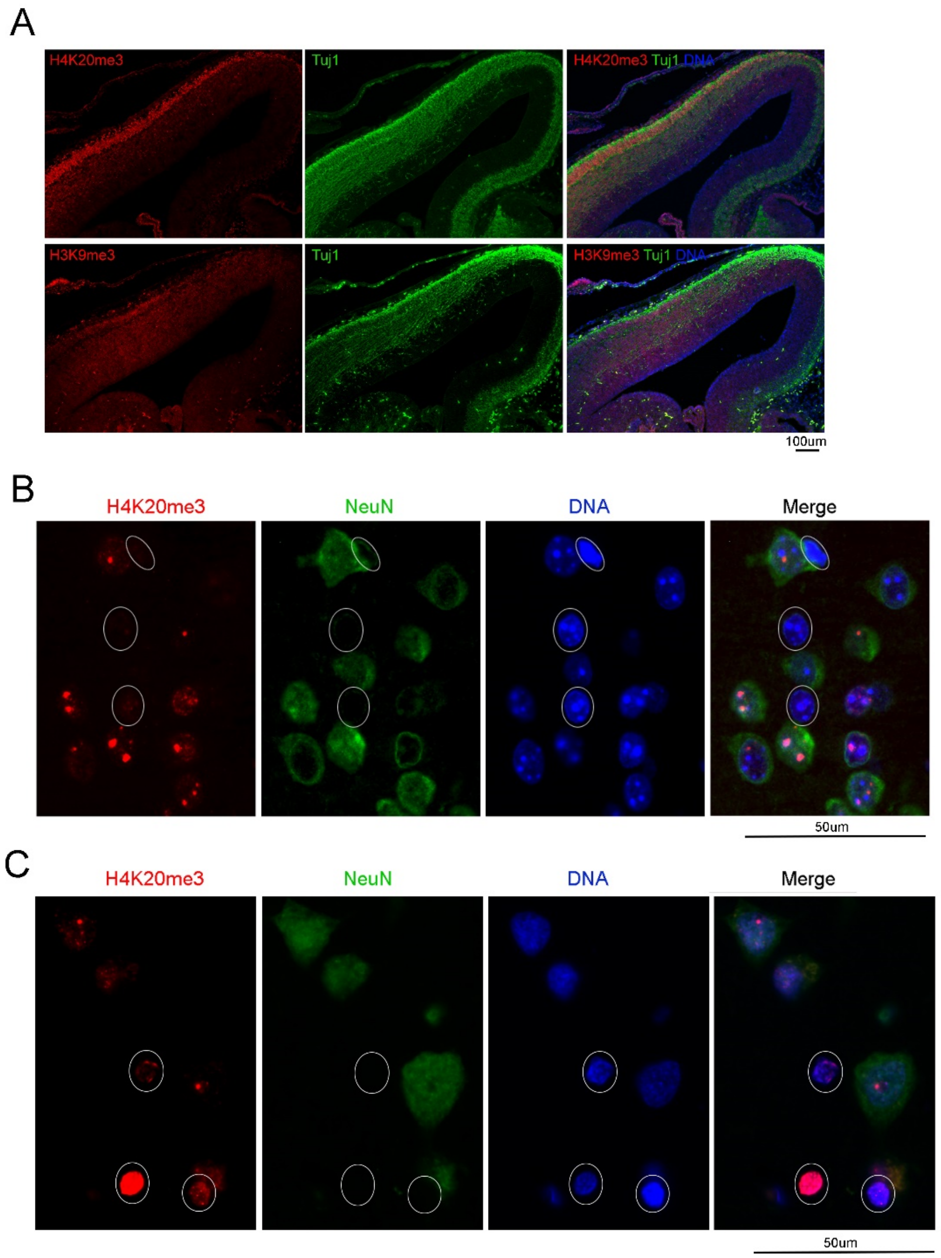


**Figure S1. H4K20me3 Modification of Heterochromatin in Neocortical Neurons**

(A) IH analysis of embryonic cerebral cortical sections with antibodies against H3K9me3 and H4K20me3, respectively. Sections were co-stained with new neuron markers Tuj1 or DCX to reveal the intense H4K20me3 foci in neurons of the nascent cortical plate at E14.5.

(B) Representative IH images of H4K20me3-NeuN double labeled cortical sections. Note the weaker H4K20me3 signals in NeuN- glia (indicated by circles).

(C) Representative IH images of H4K20me3-NeuN double labeled postmortem human cortical sections. Note the simpler H4K20me3 patterns in NeuN- glia (indicated by circles).

Nuclear DNA or chromocenters were stained by Hoechst 33342.

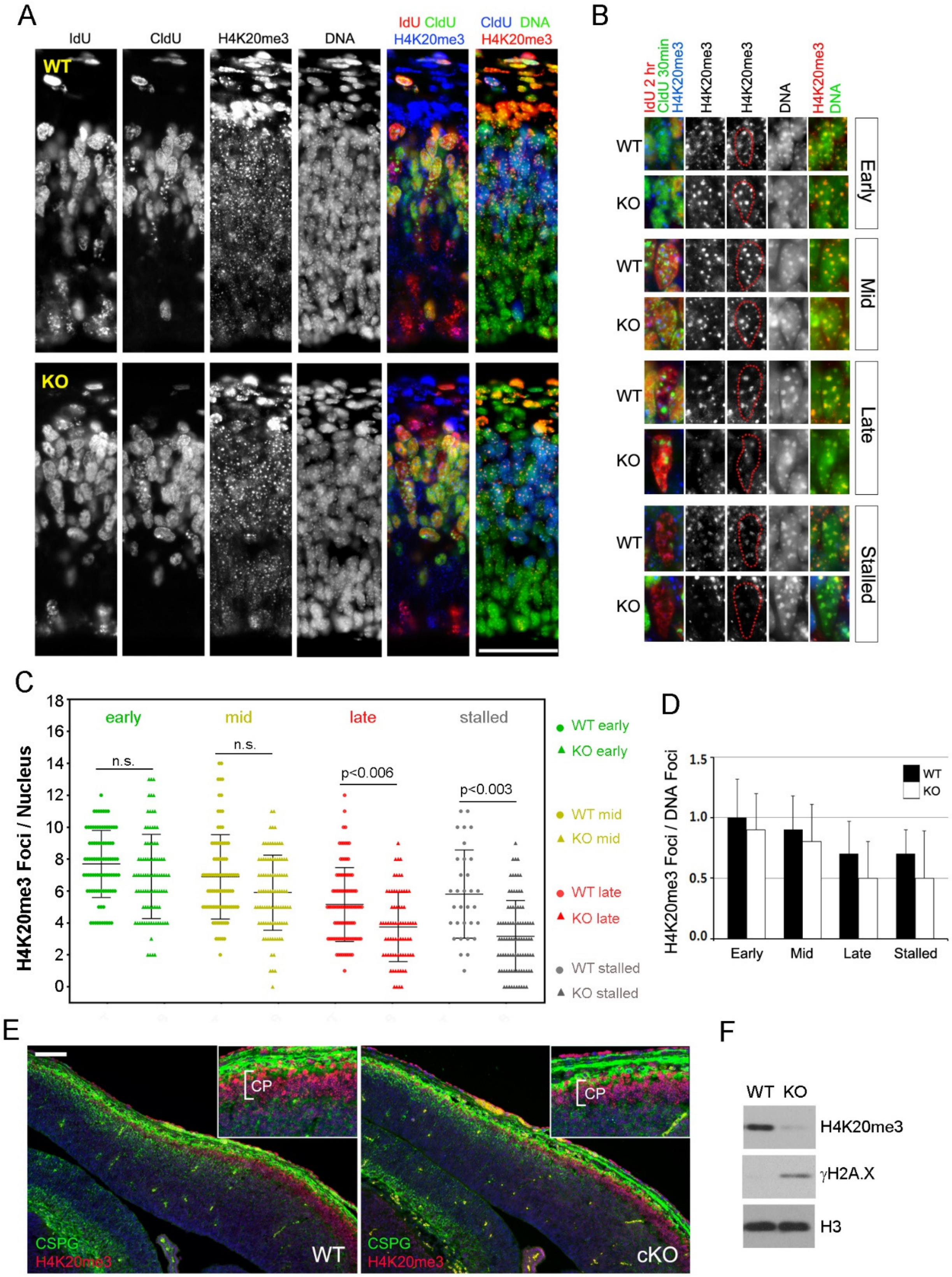
A
IdU
CldU
H4K20me3
DNA
IdU CldU H4K20me3
CldU DNA H4K20me3
WT
KO
B
IdU 2 hr CldU 30min H4K20me3
H4K20me3
H4K20me3
DNA
H4K20me3 DNA
WT
KO
Early
Mid
Late
Stalled
C
H4K20me3 Foci / Nucleus
early
mid
late
stalled
n.s.
n.s.
p<0.006
p<0.003
WT early
KO early
WT mid
KO mid
WT late
KO late
WT stalled
KO stalled
D
H4K20me3 Foci / DNA Foci
WT
KO
Early
Mid
Late
Stalled
E
CP
CP
CSPG
H4K20me3
WT
CSPG
H4K20me3
cKO
F
WT KO
H4K20me3
γH2A.X
H3

**Figure S2 Compromised H4K203 in Brap mutant cortical NPCs**

(A) Representative IH images of H4K20me3 in wild type (WT) and $Brap^{KO}$ S-phase cortical NPCs sequentially labeled by IdU (2 hours) and CldU (30 min) at E12.5. Bar: 50um.

(B) Representative IH images of H4K20me3 in early S-phase (IdU-CldU+), mid S-phase (IdU+CldU+), late S-phase (IdU+, sparse CldU+ foci), or stalled S-phase (IdU+CldU- in S-phase zone) NPCs at E12.5. H4K20me3 and Hoechst 33342 double stained images are included to reveal the H4K20me3 occupancy at chromocenters. Nuclear territory is marked by red dots on H4K20me3 images according to DNA stains by Hoechst 33342 (middle column). Note the weak and diffuse H4K20me3 immunosignals in late and stalled S-phase $Brap^{KO}$ NPCs.

(C) Quantification of the number of H4K20me3+ foci per nucleus in NPCs at early, mid, late S-phase or DNA replication was stalled in embryos of E12.5.

(D) Ratios of H4K20me3+ to Hoechst 33342+ foci in NPCs at early, mid, late S-phase or DNA replication was stalled in embryos of E12.5. Mean ± SD.

(E) IH images of H4K20me3 in control and $Brap^{cKO}$ embryos at E13.5. Note the reduced H4K20me3 in the cortical plate (CP) of $Brap^{cKO}$ embryos. Bar: 100um.

(F) IB histone extracts from cerebral cortical tissues of WT and $Brap^{KO}$ embryos.

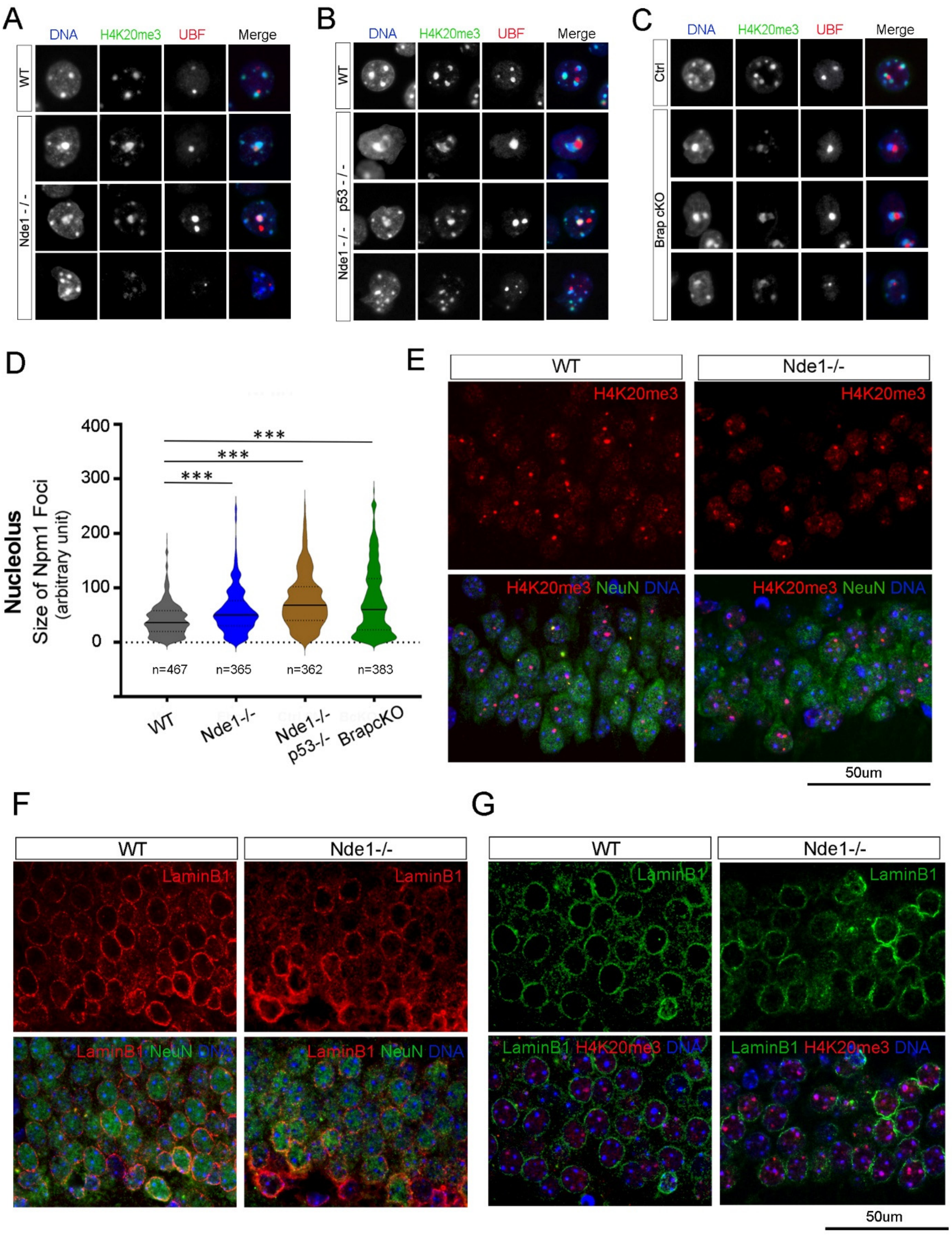
A
DNA
H4K20me3
UBF
Merge
WT
Nde1 - / -
B
WT
Nde1 - / - p53 - / -
C
Ctrl
Brap cKO
D
Nucleolus
Size of Npm1 Foci
(arbitrary unit)
400
300
200
100
0
***
n=467
n=365
n=362
n=383
WT
Nde1-/-
Nde1-/- p53-/-
BrapcKO
E
WT
Nde1-/-
H4K20me3
H4K20me3 NeuN DNA
50um
F
LaminB1
LaminB1 NeuN DNA
G
LaminB1
LaminB1 H4K20me3 DNA
50um

**Figure S3. Nde1 and Brap LOF Result in Nuclear Architecture Aberrations in Neocortical Neurons**

(A-C) IH analyses of nuclear architecture aberrations associated with nucleoli and nucleolar heterochromatin in mice of indicated genotypes. Shown are representative images of nuclei from cortical sections double stained with antibodies against UBF and H4K20me3. $n \geq 3$ for each genotype.

(D) Violin plots of nucleoli size distribution in cortical neurons of mice at weaning age (P22-28). Cortical sections were double stained with antibodies against Npm1 and NeuN. The total number of Npm1+ nucleoli quantified for each genotype ($n \geq 3$ mice per genotype) are indicated. ***$p<0.001$ by two-sample Kolmogorov-Smirnov tests.

(E) IH images of H4K20me3-NeuN double immuno-labeled brain sections of WT and $Nde1^{-/-}$ mice at weaning age (P22-28). Shown are representative images of hippocampus CA2 regions.

(F) IH images of LaminB1-NeuN double immuno-labeled brain sections of WT and $Nde1^{-/-}$ mice at weaning age (P22-28). Shown are representative images of hippocampus dentate gyrus.

(G) IH images of H4K20me3-Lamin B1 double immune-labeled brain sections of WT, $Nde1^{-/-}$ mice at weaning age (P22-28). Shown are representative images of hippocampus CA2 regions.

Bars: 50um.
